# Whole-brain cellular-resolution functional network properties of seizure susceptibility

**DOI:** 10.64898/2026.04.04.715761

**Authors:** Wei Qin, Jessica Beevis, Maya Wilde, Sarah Josephine Stednitz, Josh Arnold, Manxiu Ma, Itia Favre-Bulle, André Peterson, Ellen Hoffman, Ethan K. Scott

## Abstract

Despite its prevalence and clinical impacts, epilepsy remains incompletely understood in terms of the population dynamics that mediate seizure susceptibility, initiation, and propagation across brain-wide networks. In this study, we have performed calcium imaging in zebrafish, brain-wide and at cellular resolution, at baseline and as seizures are induced using the GABA_*A*_ receptor antagonist pentylenetetrazol (PTZ). We have then modeled the network architecture in wild-type and *scn1lab*^−/−^ larvae, which are seizure-prone and serve as a model for Dravet syndrome. *scn1lab*^−/−^ larvae show increased pair-wise correlations between neurons when exposed to PTZ, and graph analyses of these correlations revealed genotype-specific network alterations during seizures, identifying regions and metrics linked to seizure onset. Using generative network modeling, we then explored the wiring rules that govern activity in these networks, identifying specific network properties linked to seizure susceptibility that were only detectable using large-scale, cellular-resolution data. Even at baseline in the absence of seizures, these rules differed by genotype in a way that enabled the identification of *scn1lab*^−/−^ larvae and predicted individuals’ seizure risk independently of their observable phenotype. These findings uncover the cellular-resolution network properties of a zebrafish model of Dravet syndrome and establish a predictive framework for seizure susceptibility grounded in multi-scale functional connectivity.

## INTRODUCTION

Epilepsy is a neurological disorder characterized by recurrent abnormal electrical activity that results in seizures. In Dravet syndrome, both cortical and subcortical seizures are observed, originating from the cerebral cortex or deeper structures such as the thalamus^1^. These seizures arise from hyper-synchronous electrical discharges within localized brain regions, known as seizure foci, disrupting perception, behavior, and cognition^2^. Despite the availability of anti-epileptic drugs, ∼30% of patients remain pharmacoresistant^3^, facing elevated risks of sudden death, psychosocial burden, and poorer clinical outcomes^4,5^.

Given the widespread neuronal hyper-synchrony underlying seizure dynamics, characterizing whole-brain seizure networks is essential to understand ictogenesis. Previous attempts at modeling whole-brain seizure networks have addressed baseline properties that predispose networks to seizures, subthreshold events preceding seizure initiation, the spatial and temporal dynamics of seizure propagation, and pharmacological agents that can mitigate or prevent seizures^6–12^. However, traditional imaging modalities lack either cellular resolution (e.g., fMRI, EEG) or whole-brain coverage (e.g., electrophysiology), limiting the insights that can be made about multiscale network dynamics. This gap between single-cell activity and whole-brain organization prevents the modeling of seizures in a unified network framework.

Zebrafish larvae, due to their small size and optical transparency, enable whole-brain calcium imaging at cellular resolution^13^, providing a scalable platform for investigating seizure networks across multiple spatial scales. At larval stages commonly used for imaging, the zebrafish brain contains approximately 100,000 neurons^14^ and features all major mammalian neurotransmitter systems^15^. While the zebrafish telencephalon differs markedly from its mammalian counterpart, the rest of the brain, spinal cord, and core sensory and motor pathways are well conserved^16^. Seizures can be elicited in larval zebrafish through genetic manipulation and/or ictogenic drugs such as pentylenetetrazol (PTZ)^17,18^. Mutations in the zebrafish sodium channel gene *scn1lab*, homologous to mammalian *SCN1A*, recapitulate key features of epilepsy^18–21^, enabling both mechanistic studies and high-throughput drug screening^22,23^. These studies confirm that whole-brain seizure dynamics are conserved across vertebrate species^20^.

Building upon past work, we have induced seizures using PTZ in both *scn1lab*^−/−^ larvae and their wild-type (WT) siblings, and performed whole-brain calcium imaging to compare their baseline and ictogenic network dynamics. In this context, we have used PTZ as a system-wide perturbation and studied the different ways in which WT versus *scn1lab*^−/−^ networks are altered as PTZ diminishes inhibitory drive. We have found that, under baseline conditions, *scn1lab*^−/−^ larvae exhibited similar neural activity to WT. However, PTZ exposure triggered a pronounced increase in global synchrony, particularly via enhanced inter-hemispheric correlations in mid- and hindbrain neurons in *scn1lab*^−/−^ larvae, suggesting elevated contralateral functional connectivity during pre-seizure states. Graph theory analyses revealed distinct PTZ-driven connectivity patterns in *scn1lab*^−/−^ brains, with early divergence in regions such as the pallium and habenula.

We then used generative network modeling (GNM), which analyzes the statistical properties of network architecture to infer connectivity principles, and revealed that baseline network topology alone could classify genotype and predict seizure susceptibility. Specifically, we found that *scn1lab*^−/−^ networks are defined by distinct generative parameters, where unique mathematical wiring principles deviate from wild-type norms. This suggests latent organizational vulnerabilities that predispose *scn1lab*^−/−^ larvae to hyper-synchronization. These findings advance our understanding of multi-scale seizure network dynamics and offer a framework for predictive modeling and therapeutic targeting.

## RESULTS

### Increased susceptibility to PTZ-induced seizures in *scn1lab*^−/−^ animals

To monitor neuronal activity and movement simultaneously during seizure induction, we developed a custom imaging chamber coupled with a dual-objective light-sheet fluorescence microscope (Figure 1A; see Online Methods). Larval zebrafish were embedded in low-melting-point agarose with their heads immobilized and tails free to move. Volumetric calcium imaging was achieved by illuminating the brains of elavl3:H2B-GCaMP6s larvae (expressing the calcium indicator GCaMP6s in the nuclei of all neurons^24^) with two orthogonally oriented light sheets, one anterior and one lateral, while collecting fluorescence signals through an objective positioned above the animal^13,25^. Concurrently, tail movements were recorded using infrared LEDs, a low-power objective beneath the chamber, and a high-speed camera (see Online Methods). All larvae were progeny of in-crossed *scn1lab*^+/−^ adults and were genotyped post hoc.

**Figure 1.**
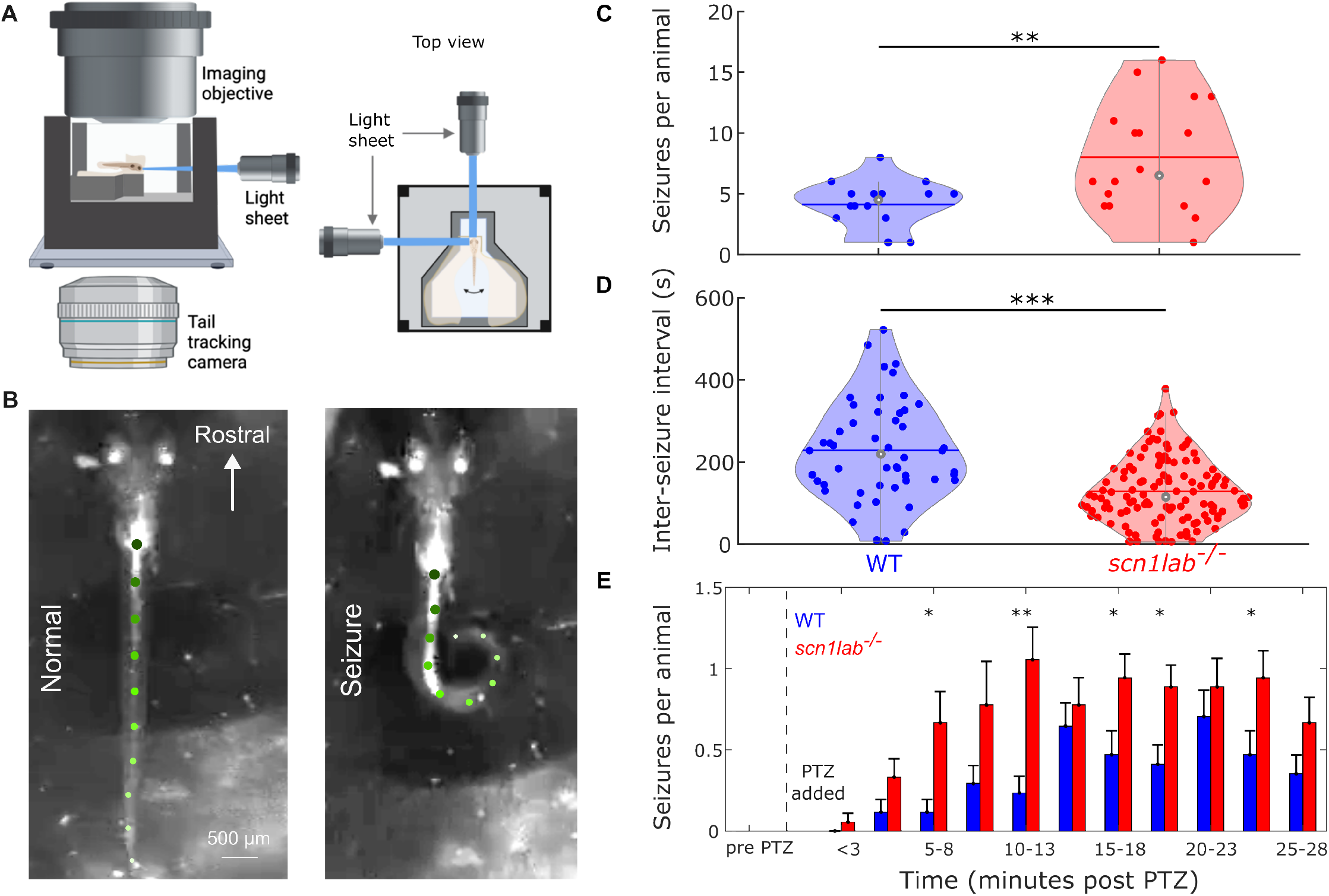
Experimental setup and seizure susceptibility. A custom whole-brain imaging and tail-tracking setup reveals that *scn1lab*^−/−^ mutant larvae exhibit significantly higher seizure frequency and shorter inter-seizure intervals than WT siblings following PTZ administration, confirming increased seizure susceptibility. (A) A schematic of the experimental preparation. (Left) A custom chamber holds the animal’s head still on a stage. An orthogonal imaging objective (above) allows for calcium imaging in the brain, while a second objective (below) leads to a camera for tail tracking. (Right) Top view of the imaging chamber showing the two light sheets used for calcium imaging, one from the front of the fish and one from the side. Created with BioRender.com (B) Examples of images from the tail tracking camera, with the tail’s automatically detected position indicated (green dots). The tail is nearly straight and displays small, slow movements between seizures (Left), but exhibits dramatic movements and sustained high-amplitude bends during seizures (Right). The original recording of the seizure event is presented in Video 1. (C) The average numbers of seizures shown by *scn1lab*^−/−^ (red) and WT sibling (blue) larvae during the course of the experiment (n=17 and 18, respectively, ** = p *<* 0.01, Student’s T-test). (D) The inter-seizure intervals for the two genotypes, showing that seizures occur more frequently in *scn1lab*^−/−^ animals (148 seizures across 35 animals, *** = p *<* 0.001, Student’s T-test). (E) Mean seizure counts per genotype (*scn1lab*^−/−^ red; WT blue) are plotted across time following PTZ administration (n = 17 and 18). Error bars represent the standard error of the mean (SEM). Statistical significance was determined by analysis of variance (ANOVA) with Šidák’s correction (* = p *<* 0.05, ** = p *<* 0.01).

Following a five-minute baseline observation of brain activity and tail movements, we applied 5 mM PTZ to induce seizures. This approach was based on previous studies showing that *scn1lab*^−/−^ larvae exhibit a lower threshold for PTZ-induced seizures^17,26^. Consistent with previous studies, we observed episodes of rapid, high-frequency tail flicking and convulsive movements^26^ (Figure 1B). In our preparation, neither *scn1lab*^−/−^ nor WT animals showed spontaneous seizures. Indeed, in free-swimming experiments, we were unable to induce seizures in *scn1lab*^−/−^ larvae even with intense acoustic^27^, visual^28^, or combined stimuli, suggesting that these animals are not susceptible to seizures in the absence of PTZ, and that they have, in our hands, a latent predisposition for seizures.

Volumetric whole-brain imaging was performed at 2 Hz with a 10 *µm* step along the dorsalventral axis, generating 25 planes per animal. For each animal, the recorded image stack was processed using Suite2p to identify regions of interest (ROIs) corresponding to individual neurons, and to extract the calcium activity trace associated with each neuron^29^. The same image stacks were subsequently registered to the Z-Brain atlas using ANTs^30,31^. Finally, the extracted cellular activity was mapped to anatomical regions within Z-Brain space using custom-written MATLAB scripts^32,33^ (Figure 2A). We applied four behavioral or neural activity criteria to identify seizure episodes (see Online Methods): mean activity of all neurons across the brain (mean Δ*F/F*_0_ beyond 99% interquartile range, Figure 2C), synchronization of this activity (skewness of the network correlation distribution *>* 0.8, Figure S1), movements of the head as measured by motion correction (frame-to-frame lateral motion dynamics extracted from Suite2p beyond 99% interquartile range, see Online Methods), and tail movements (mean tail angle beyond 99.9% interquartile range, Figure 2C) ^17,22,34,35^. We found that PTZ-induced seizures occur earlier, more frequently, and in greater numbers in *scn1lab*^−/−^ animals than in their WT siblings (Figure 1C-E, S1E).

**Figure 2.**
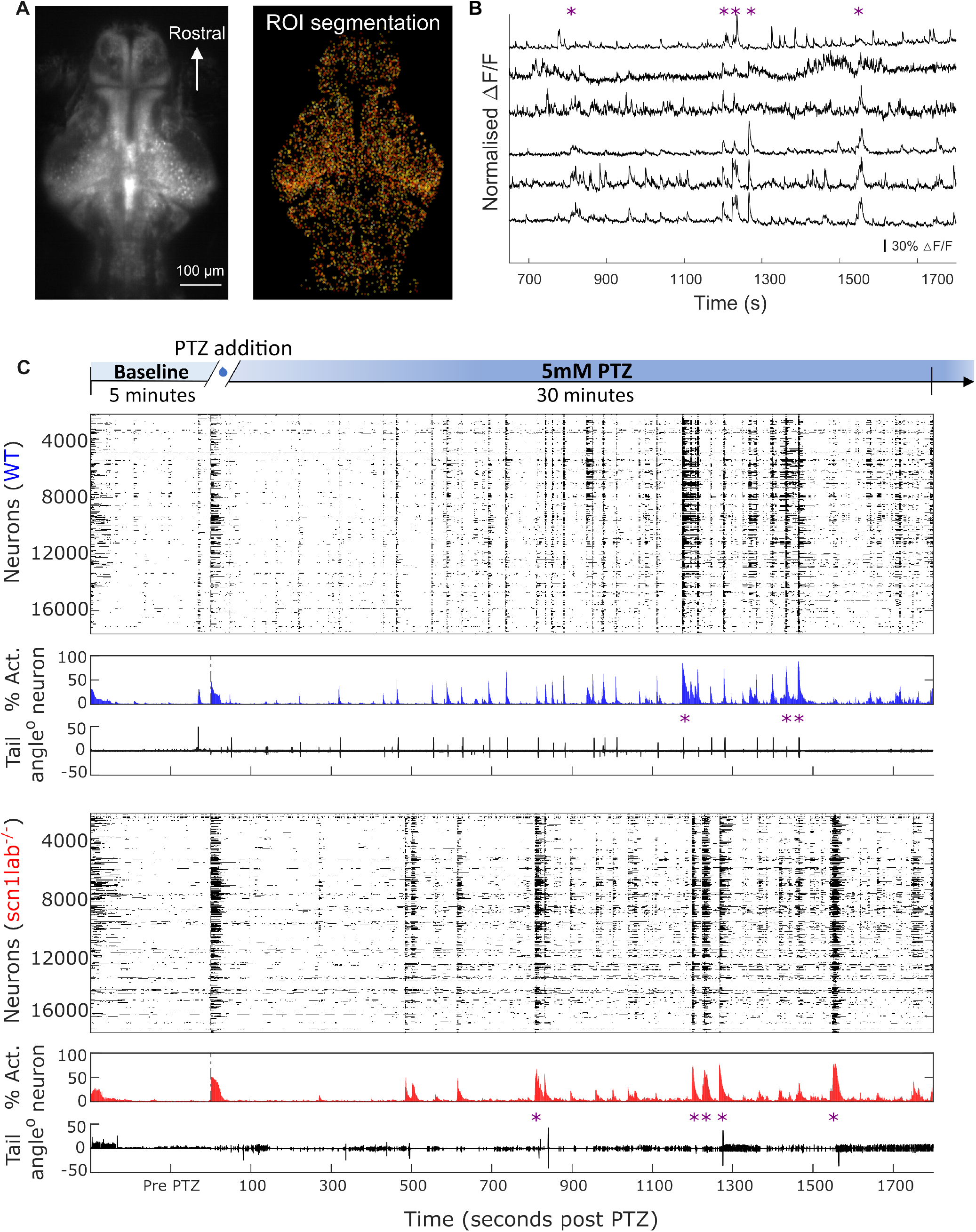
Whole-brain imaging of seizure activity at cellular resolution. Whole-brain calcium imaging at cellular resolution captures distinct seizure dynamics in WT and *scn1lab*^−/−^ mutant larvae, revealing genotype-specific differences in neuronal activations and seizure patterns across the PTZ experiment. (A) A maximum intensity projection of a brain during our calcium imaging experiment (Left) and the locations of the neurons segmented from these raw data (Right). (B) Example traces of seven neurons from this dataset, showing each neuron’s activity during a portion of the experiment. Seizure events are indicated with an asterisk. (C) Raster plots of activity across all neurons for one WT animal (blue, top) and one *scn1lab*^−/−^ animal (bottom, red), across the entire experiment (thresholding criteria are described in Online Methods). The percentage of active neurons across the brain and tail movements are also shown for the duration of the experiment. Figure S1 provides detailed lists of mean calcium activity for each individual fish. Seizure events (three for the WT and five for the *scn1lab*^−/−^ animal) are indicated with asterisks.

### The number, density, and activity of neurons are altered across the *scn1lab*^−/−^ brain

We next explored the functional properties of the seizure networks using the activity profile of each neuron (see Figures 2B-C). The cell coordinates, representing the center of mass of each Suite2p-generated ROI, were carried forward and converted into their respective atlas positions (see Online Methods). To assess regional differences in neuronal recruitment, we segmented the brain into four major anatomical domains: the telencephalon and diencephalon (collectively the forebrain), mesencephalon (midbrain), and rhombencephalon (hindbrain) (Figure 3A). Using the extracted cellular coordinates for registered neurons, we then analyzed morphological changes in brain regions between genotypes, applying both the ROI density method (comparing the number of neurons found within prescribed brain volumes) and a previously published Symmetric Normalization (SyN) method (measuring local volume changes across spatially registered brain images^36^, see Online Methods). Our findings revealed significant differences in ROI densities (*ω*) across various brain areas (Figure 3B; A complete stack of planes across the dorsoventral axis is provided in Figure S2). Proportionally more ROIs were segmented in the forebrain in WT larvae, whereas *scn1lab*^−/−^ had more in the midbrain and hindbrain (Figure 3C). Despite these regional shifts, neither the proportional nor absolute number of ROIs differed significantly across major brain divisions. Total ROI counts across the whole brain were also indistinguishable between genotypes (Figure S1D).

**Figure 3.**
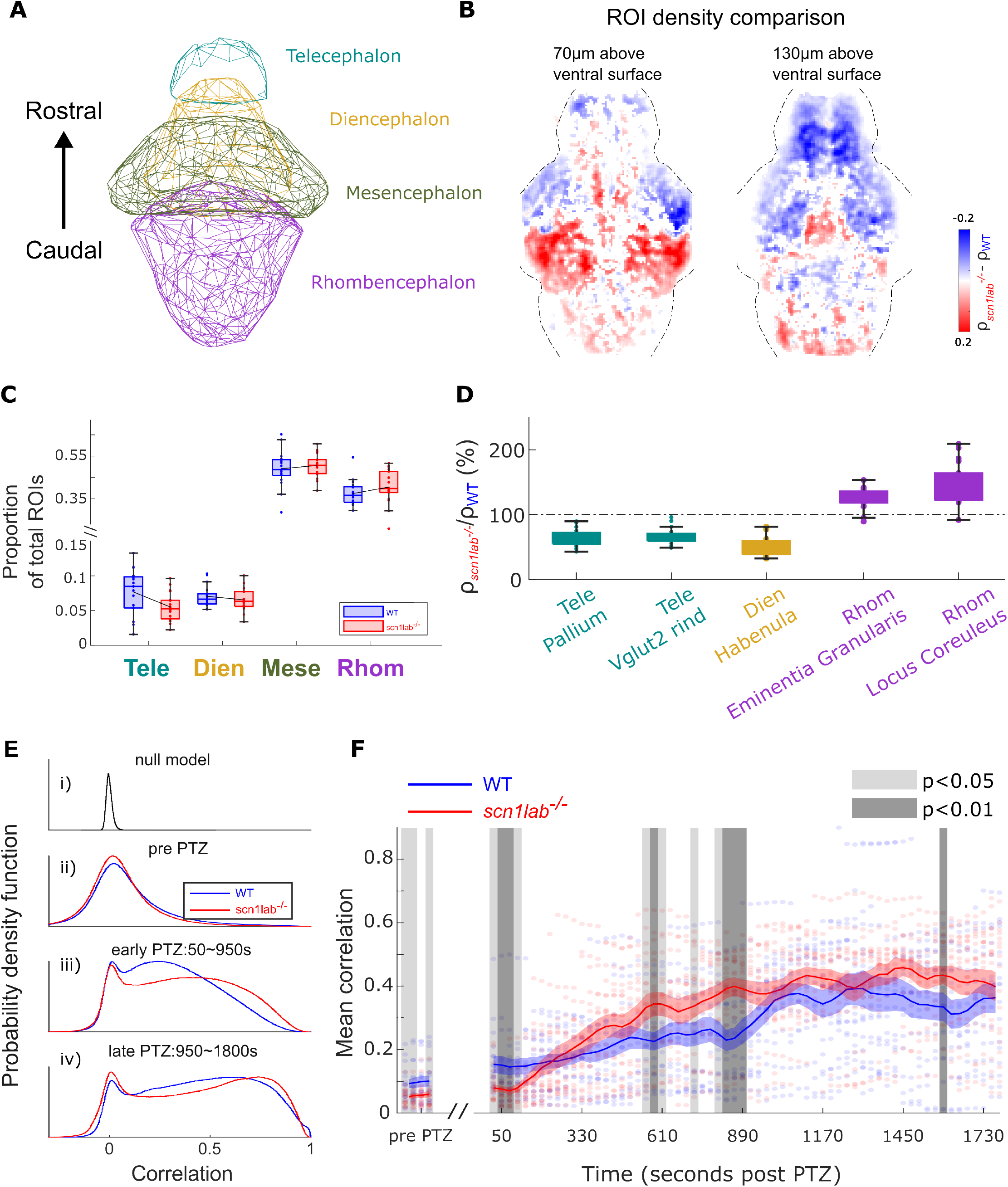
Spatial and functional alterations in neurons across *scn1lab*^−/−^ brains. *scn1lab*^−/−^ mutant brains exhibit region-specific changes in neuronal density and altered functional connectivity dynamics, revealing genotype-dependent spatial and temporal reorganization of neural activity. (A) Anatomical segmentation of the brain into four major regions: telencephalon, diencephalon, mesencephalon, and rhombencephalon. (B) A voxel-based analysis of ROI density (*ρ*) across *scn1lab*^−/−^ and WT brains. Two different horizontal planes are shown, with depth above the ventral surface indicated. Blue voxels, mostly in the forebrain, have significantly greater density and number of neurons in WT animals, and red voxels, with higher numbers and densities in *scn1lab*^−/−^ larvae, are concentrated in the hindbrain. Voxels are colored if they have p *<* 0.05 (Mann-Whitney U test). ROI distribution comparisons between genotypes: (C) ROI number proportions across major brain regions, and (D) Subregion-level neuron ratios with p *<* 0.01. Statistical significance was assessed using the Mann-Whitney U test. Figure S2 displays all subregions showing significant differences at p *<* 0.05. (E) Correlation distributions (null vs. real, pre- and post-PTZ) across the course of the experiment. (F) Correlation shifts over time. Each dot represents one animal’s mean Pearson’s correlation. Blue and red denote WT and *scn1lab*^−/−^ genotypes, respectively. Solid lines indicate group means, with shaded regions representing the SEM. Gray shading marks timepoints with significant genotype effects, as determined by repeated-measures ANOVA with Bonferroni correction. Two significance thresholds are visualized by differential shading intensity.

To resolve finer spatial patterns, we extended this analysis to 137 anatomically defined subregions of the brain. ROI counts were normalized to mean WT values, and subregions shortlisted with significant genotype-dependent differences (p *<* 0.01, Mann—Whitney U test) were identified (Figure 3D). Consistent with our observations of broad regions, *scn1lab*^−/−^ larvae exhibited reduced ROI counts in forebrain subregions, including the pallium and habenula, alongside increased numbers in hindbrain areas such as the eminentia granularis (EG) and locus coeruleus. A complete list of subregions with p *<* 0.05 (Mann-Whitney U test) is provided in Figure S2.

Using our whole-brain, cellular-resolution activity data, we next examined network-wide functional correlations across genotypes and timepoints. For each pair of neurons in an individual brain, Pearson’s correlation coefficients were calculated. A time-shuffled null model preserved each animal’s activity distribution while eliminating temporal structure, yielding a narrow distribution centered near zero (Figure 3Ei). In contrast, unshuffled activity data revealed broader, positively skewed correlations at baseline, with minimal differences between *scn1lab*^−/−^ and WT larvae before PTZ (Figure 3Eii), suggesting that baseline synchrony does not predispose *scn1lab*^−/−^ animals to seizures.

Upon PTZ application, correlation distributions shifted significantly in the positive direction within the first 15 minutes, with *scn1lab*^−/−^ larvae showing a stronger shift than WT (Figures 3Eiii and 3F). During the subsequent 15-minute interval, WT larvae showed continued increases in correlation strength, ultimately reaching levels comparable to *scn1lab*^−/−^ larvae by the end of the experiment (Figure 3Eiv). These shifts in correlation coincided with seizure onset and progression, which occurred earlier and more robustly in *scn1lab*^−/−^ animals. WT larvae displayed delayed and less consistent seizure dynamics, but ultimately converged toward similar network states. These findings implicate elevated pairwise neuronal correlations as a network-level marker of ictogenesis.

### PTZ drives a dramatic increase in inter-hemispheric connectivity in *scn1lab*^−/−^ animals

To determine whether these correlation changes were spatially uniform or modulated by anatomical and spatial factors, we examined the influence of brain region, intercellular distance, and hemispheric pairing. Consistent with prior studies, pairwise correlations generally declined with increasing distance between neurons (Figure 4A)^37,38^. Similar to Figure 3F, Figure 4A demonstrates elevated mean of whole-brain correlations across all inter-neuronal distances at baseline. Following PTZ administration, *scn1lab*^−/−^ larvae exhibit a pronounced increase in functional connectivity (defined as correlation strength), which gradually converges with WT during the late PTZ period, coinciding with frequent seizure activity. Interestingly, *scn1lab*^−/−^ larvae also exhibited an unexpected rise in correlation for neuron pairs separated by 300–450 *µm* (Figure 4B), a range typically corresponding to contralateral hemispheric pairings^39^. Genotype-specific differences in both power coefficients and model fit imply altered spatial wiring principles in *scn1lab*^−/−^ brains, as determined by linear regression analysis (Figure S3).

**Figure 4.**
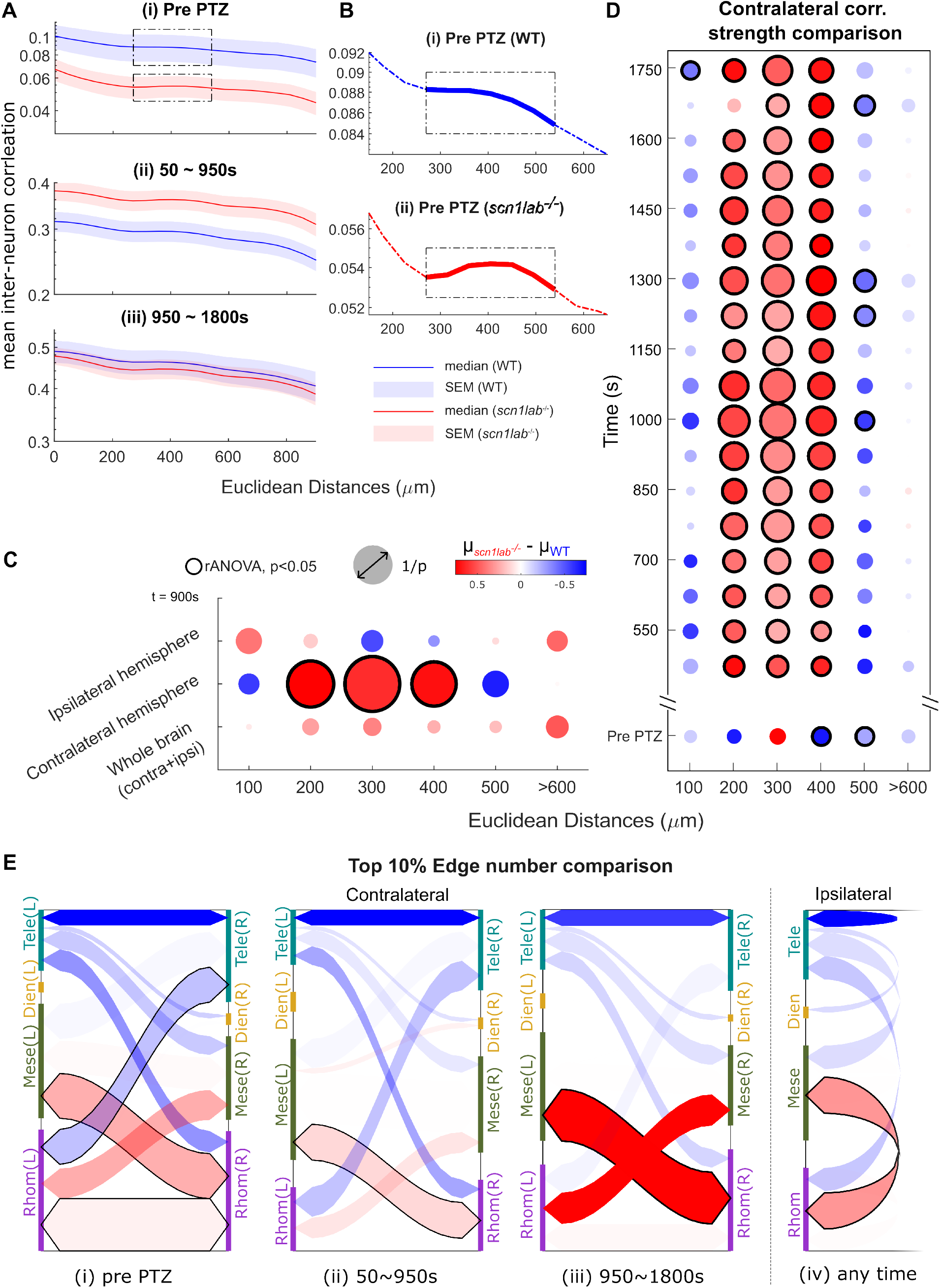
Hemispheric functional connectivity is altered for *scn1lab*^−/−^ larvae in PTZ. *scn1lab*^−/−^ mutant larvae exhibit altered hemispheric connectivity under PTZ, with enhanced contralateral correlations and hindbrain enrichment that diverge from WT patterns, revealing genotype-specific reorganization of long-range functional interactions. (A) Correlation strength as a function of neuron distance during pre-(i), early (ii), and late (iii) PTZ stages. WT and *scn1lab*^−/−^ larvae show declining correlations with increasing distance. (B) Close-ups of correlations within the boxes in Ai, showing a range in which correlations increase with Euclidean distance in *scn1lab*^−/−^ animals. (C) Correlation strength 900 s after PTZ application for ipsilateral, contralateral, and whole-brain pairings at different Euclidean distances. Blue: WT correlations are greater than *scn1lab*^−/−^, red: *scn1lab*^−/−^ correlations are greater than WT, and black circles: p *<* 0.05 (rANOVA, with Bonferroni Correction). (D) Contralateral correlation strengths are stronger in *scn1lab*^−/−^ throughout the PTZ period for pairings between 200-400 *µm*. Y axis indicates the time points since PTZ administration. Detailed comparisons of ipsilateral and contralateral regional and subregional connectivity are presented in Figure S3. (E) Top 10% of pairwise correlations mapped to anatomical regions. Contralateral connections over three PTZ stages (i) pre-PTZ, (ii) 50 ∼ 950 s and (iii) 950 ∼ 1800 s post PTZ. *scn1lab*^−/−^ larvae show enrichment in hindbrain connectivity. Ipsilateral connections remain unchanged as in (iv). Arrow color denotes mean differences; width reflects statistical strength (1/p from repeated-measures ANOVA with Bonferroni correction). Significant connections are outlined in black.

We next explored the importance of ipsilateral versus contralateral connectivity across genotypes. During PTZ exposure, contralateral neuron pairs in *scn1lab*^−/−^ larvae showed markedly elevated correlation strengths within the 200–400 *µm* range, exceeding those observed in WT siblings (Figure 4C). This effect was prominent during PTZ exposure, but was also weakly present at baseline for contralateral pairs near 300 *µm* in *scn1lab*^−/−^ larvae (Figure 4D), contrasting with the higher baseline whole-brain synchrony observed in WT animals (Figures 3E and 4A). A region-wise comparison of correlation strengths across major brain divisions and their subregions is provided in Figure S3.

To dissect the spatial distribution of high-correlation pairings, we normalized across genotypes and mapped the top 10% of pairwise correlations across each brain (Figure 4E). At baseline, the highest-correlation connections in *scn1lab*^−/−^ larvae were concentrated within the rhombencephalon and between the rhombencephalon and mesencephalon, whereas WT siblings had more spatially distributed high-correlation pairings across other brain regions. A detailed comparison of each major region with its respective subregions is provided in Figure S3.

Together, these findings imply that elevated hindbrain synchrony and reduced forebrain correlations in *scn1lab*^−/−^ larvae may contribute to their heightened susceptibility to PTZ-induced seizures. These patterns implicate strengthened inter-hemispheric functional connectivity as a key contributor to seizure dynamics in this model.

### Graph theory analysis reveals altered functional network structure across *scn1lab*^−/−^ brains

To move beyond pairwise correlations and assess higher-order functional organization, we constructed brain-wide networks where nodes represent individual neurons and edges represent functional connectivity (correlation exceeding a defined threshold). We then applied a suite of eight graph theory metrics (Table 2) drawn from the Brain Connectivity Toolbox^40^ to these networks to identify phenotypic differences between *scn1lab*^−/−^ and WT larvae.

We found numerous ways in which the genotypes differed across these metrics in their responses to PTZ (shown in Figure 5A). For example, assortativity is a network-wise metric that measures the tendency of nodes to connect with other nodes of similar degrees. Before PTZ, *scn1lab*^−/−^ animals have higher assortativity, indicating a greater tendency for connections between similar nodes. This observation aligns with previous studies showing that during patho-physiological conditions, networks often have high assortativity, characterized by a resilient core of highly interconnected nodes that facilitates synchronization^41–44^. As shown in Figure 5A, assortativity effectively differentiates the genotypes before PTZ application, but this difference across genotypes is reduced after PTZ exposure.

**Figure 5.**
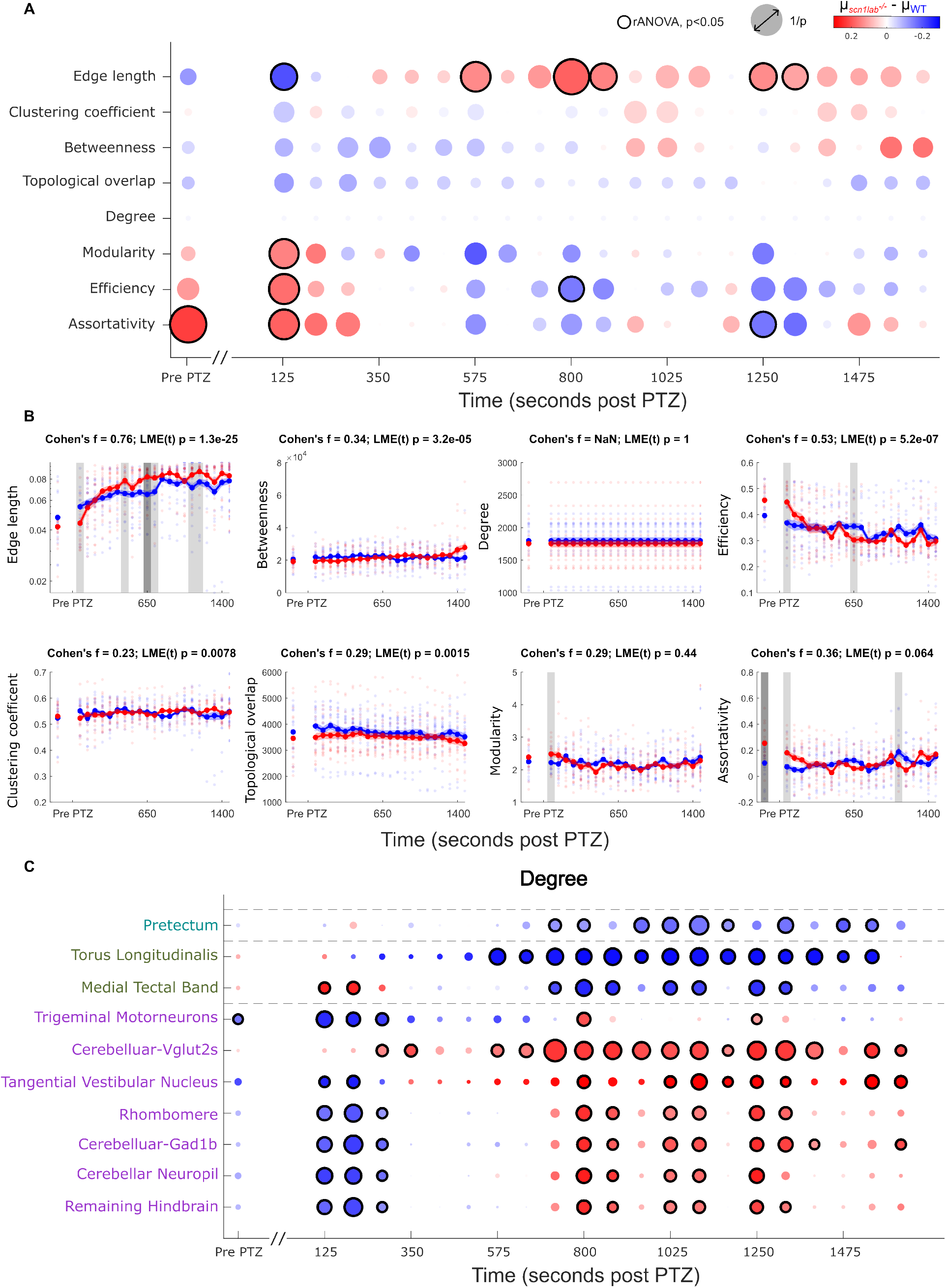
Graph metrics across WT and *scn1lab*^−/−^ brains. Graph-theoretical analysis reveals genotype-specific alterations in network property over time, with *scn1lab*^−/−^ mutants showing distinct alterations in global metrics and subregional node degree, highlighting dynamic reorganization of brain connectivity during PTZ exposure. (A) Comparison of genotype-specific changes in eight graph network metrics from pre-PTZ to post-PTZ. Blue circles indicate that the WT mean is greater and a red circle indicates that the *scn1lab*^−/−^ mean is greater. Circles outlined in black denote statistically significant differences (p *<* 0.05), determined by repeated-measures ANOVA with Bonferroni correction. (B) Time-resolved profiles of eight graph theory metrics are shown, with each dot representing the mean value for an individual animal. Blue and red denote WT and *scn1lab*^−/−^ genotypes, respectively. Solid lines indicate group means, with shaded regions representing the SEM. Gray shading marks timepoints with significant genotype effects, as determined by repeated-measures ANOVA with Bonferroni correction (p *<* 0.05). Cohen’s *f* values are reported on each panel to quantify effect size between genotypes. P-values for each network metric were obtained from LME, which assessed the statistical significance of temporal trajectories across the cohort. Summary table is shown in panel (A). (C) Of the 137 regions analyzed, only ten exhibited statistically significant differences in node degree over time. Blue circles indicate that the WT mean is greater and a red circle indicates that the *scn1lab*^−/−^ mean is greater. Circles outlined in black denote statistically significant differences (p *<* 0.01), determined by repeated-measures ANOVA with Bonferroni correction. Subregional comparisons via other metrics are shown in Figure S4.

Besides assortativity, edge length also demonstrated significant genotype-dependent differences, initially presenting higher in WT at 125 s before transitioning to significantly higher values in *scn1lab*^−/−^ during later seizure stages (e.g., 575 s and 800 s). In contrast, other metrics such as the clustering coefficient, which quantifies local interconnectedness, showed no statistically significant differences in multiple comparison analyses. However, a genotype-dependent shift (blue to red) was observed during seizures, with mean values transitioning from higher in WT to higher in *scn1lab*^−/−^. Similar trends were noted for betweenness centrality. Modularity, reflecting the extent of community structure, and network efficiency displayed comparable patterns, with marginal significance in select post-PTZ intervals. Notably, efficiency metrics demonstrated an inverse shift relative to clustering, transitioning from higher values in *scn1lab*^−/−^ to WT.

Node-level metrics were computed by averaging across all nodes (see Online Methods). Topological overlap, a measure of shared connectivity between neurons, was marginally higher in WT networks across the experimental timeline. Betweenness centrality, which captures the extent to which nodes mediate shortest paths, mirrored the clustering coefficient trend, shifting from higher WT to higher *scn1lab*^−/−^ values post-PTZ. Node degree remained stable across conditions, reflecting the fixed binarization threshold (10%) used in network construction. In contrast, edge length emerged as a robust genotype-sensitive metric, particularly during seizure-intensive epochs. During these periods, *scn1lab*^−/−^ larvae exhibited significantly greater edge lengths compared to WT controls (indicated by the large red bubbles). This sustained divergence suggests that the mutant network undergoes a distinct form of PTZ-induced reorganization.

Figure 5B summarizes the eight metrics’ temporal dynamics across genotypes. Following PTZ administration, edge length and betweenness centrality exhibited strong increases, accompanied by a corresponding decline in global efficiency, indicative of impaired signal (since it is inversely proportional to the path length). These longitudinal trends were statistically robust, with highly significant time effects in the Linear mixed-effects model (LME p *<* 0.001; see Online Methods), and were further supported by large genotype-specific effect sizes (Cohen’s *f* = 0.76, 0.34, and 0.53). Modularity and assortativity, metrics reflecting global network organization, displayed transient fluctuations, potentially linked to seizure episodes. Although their LME-derived p-values were high (p *>* 0.05), the large effect sizes (Cohen’s *f* = 0.29 and 0.36) suggest notable genotype-specific divergence. Additional metrics, including clustering coefficient and topological overlap, showed significant temporal changes, with low p-values (p *<* 0.01) and moderate effect sizes (Cohen’s *f* = 0.23 and 0.29), pointing to subtle genotype-dependent modulation. As expected, node degree remained stable across time points, consistent with the fixed thresholding approach used to construct binarized networks (top 10% Pearson correlation). Collectively, the temporal profiles and genotype comparisons of all metrics are presented in Figure 5B, with corresponding statistical details provided in Online Methods.

To identify genotype-dependent network features, we first applied a 10% correlation threshold and found that specific metrics, such as assortativity, reliably distinguished genotypes, whereas others did not (Figure 5A). To ensure these findings were robust to threshold selection, we performed a sensitivity analysis across a range of thresholds (5%–25%), which confirmed the consistency of the results that we found with a 10% correlation threshold (Figure S4B). We then questioned whether the observed phenotypic differences merely reflected trivial aspects of the network’s degree distribution. Comparison with Maslov–Sneppen (MS) randomized networks revealed that graph metrics significantly diverged from random expectations (Figure S4E), indicating that these measures capture biologically meaningful changes in network organization. Notably, genotype-specific differences were primarily driven by systematic variation in node degree, the number of functional connections per neuron (Figure S4C and D). This core difference in node degree likely explains broader network variations between genotypes.

Guided by this result, we performed a fine-grained analysis of degree distributions across 137 brain subregions. This subregional resolution revealed that genotypic differences were not uniform but were concentrated in specific areas, including the pretectum, torus longitudinalis, and cerebellum (Figure 5C). Notably, these differences were absent prior to PTZ administration, suggesting that the network properties predisposing animals to seizures are not made evident by measurements of baseline degree.

We extended this subregional analysis to other node-wise metrics, including betweenness centrality, edge length, and topological overlap (Figure S4A). The topological overlap analysis identifies several key subregions, such as torus longitudinalis and cerebellar granule cells, where WT exhibits significantly greater values compared to *scn1lab*^−/−^. More importantly, clear transitions occur: during pre-PTZ, WT and *scn1lab*^−/−^ properties are very similar, but this shifts to higher WT values in early PTZ, followed by higher *scn1lab*^−/−^ values in late PTZ (Figure S4Aiii). A similar trend is observed in degree (Figure 5C) and edge length (Figure S4Aii), indicating that these time-dependent changes may be closely related to network efficiency. Betweenness centrality highlights critical hubs (subregions) that play an important role in integrating activity across functionally distinct regions. WT generally exhibits higher betweenness centrality, implying higher network resilience and a more hierarchical organization (Figure S4Ai).

Collectively, these graph analyses reinforce the idea that, while subtle differences may exist across the genotypes’ functional network structures at baseline, the *scn1lab*^−/−^ animals’ susceptibility to seizures rests more on differential responses to PTZ, at least insofar as graph analyses can identify them. That being said, one significantly altered baseline measure, assortativity, suggests that *scn1lab*^−/−^ networks may have subtle baseline alterations toward interconnectedness and more robust signal propagation.

### Generative network modeling reveals the regional structure of seizure risk

Despite extensive work in humans and animal models using measures of neuronal activity, network synchrony, and graph characteristics, the underlying mechanisms of seizure susceptibility remain poorly understood. Given the hints provided by our graph analyses of whole-brain cellular-resolution activity data, we next turned to generative network modeling (GNM) to explore the altered rules by which predisposed networks function.

GNM offers a mathematical framework for understanding the formation process of a network from its inception^45^. Rather than relying on the assumption of random connectivity, GNM generates synthetic networks whose statistical and structural properties mirror those observed in empirical data. The model operates on the premise that connections between nodes follow specific wiring principles, enabling biologically plausible inference of network topology. This hypothesis-driven approach simulates the balance between heuristic reward functions and cost functions. The model calculates the pairwise connection probabilities between each pair of nodes in a network and constructs the network according to these calculated probabilities (See Online Methods).

We adapted the two-parameter growth-based generative model proposed by Betzel et al. ^46^, and inspired by the economical clustering and economical preferential attachment models^47^. We began with an unconnected network and added connections incrementally to simulate network growth. At each iteration, we introduced a new edge between nodes *i* and *j* based on the following probability:

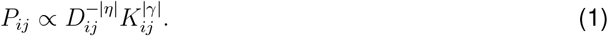

Here, *D*_*ij*_ represents the Euclidean distance between nodes *i* and *j*, while *K*_*ij*_ denotes a topological similarity that evolves as the network grows. We implemented a single homophily-based wiring rule, the matching index, as the reward function *K*. This rule quantifies the proportion of shared neighbors between two nodes, serving as a measure of local connectivity overlap. A high matching index indicates that nodes occupy similar network neighborhoods, even in the absence of direct connections. Prior work has shown that matching index is among the most effective homophily rules for recapitulating empirical brain network structure^45^. Because the matching index *K*_*ij*_ depends on the evolving network topology, it is dynamically updated with each newly formed connection. To prevent duplicate connections between the same node pair, *P*_*ij*_ is set to zero once a connection is established. We repeated this iterative process until the simulated network matched the empirical functional connectome in edge count, corresponding to the top 10% of strongest correlations in the empirical network^48^.

In this model, *η* and *γ* serve as scaling parameters that modulate the influence of spatial and topological constraints, respectively. − | *η* | penalizes long-range connections based on Euclidean distance, reflecting biological costs of wiring.|*γ* | amplifies the contribution of the reward function, the matching index, favoring connections between nodes with similar local connectivity. When *η* = 0 or *γ* = 0, the model lacks spatial or topological constraints, resulting in unregulated, random edge formation without biologically informed wiring rules. Together, these parameters allow flexible tuning of the model to balance biological cost and homophily in reproducing realistic network architecture^45^.

To generate a whole-brain coarse-grained model using this approach, individual neurons were clustered to 500 nodes per brain, using K-means, to reduce computational complexity (Figure 6A), and 300 simulations were run for all combinations of *η* and *γ* values across a broad range for each (Figure 6B). Each simulation was evaluated by comparing the Kolmogorov–Smirnov (KS) energy similarity, and by averaging the results of these simulations, we generated “fingerprints” of the 2D area of *η* and *γ* values, which reflect the values that best matched our experimental data (Figure 6C, Online Methods). Across the duration of the simulations, *η* and *γ* values are essentially stable and show no meaningful differences between the genotypes (Figures 6D-E), indicating that GNM does not reveal the root causes of seizure risk using the whole-brain coarse-grained data.

**Figure 6.**
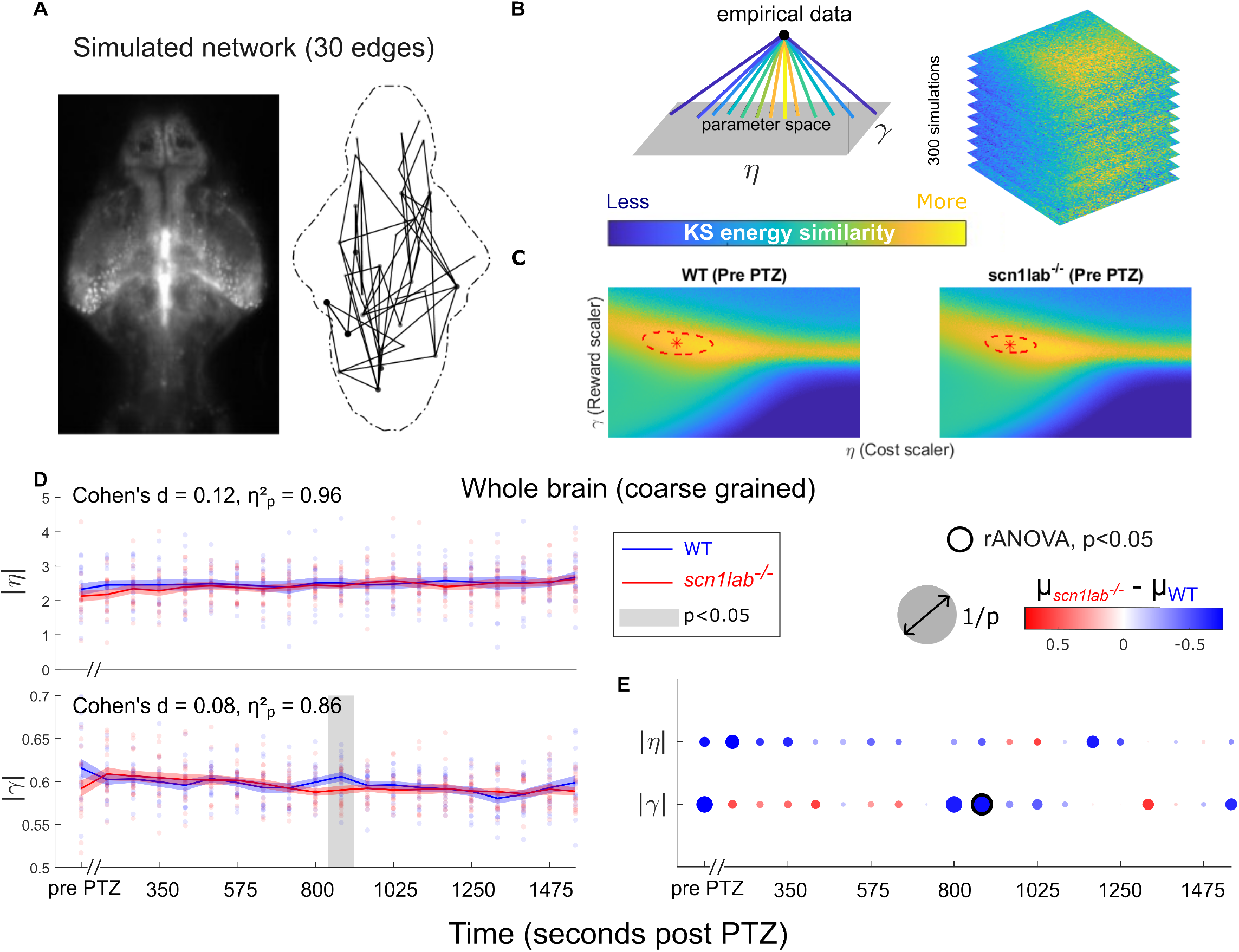
Generative network modeling of WT and *scn1lab*^−/−^ brains. Generative modeling reveals that coarse-grained wiring parameters show minimal genotype-specific differences, with *scn1lab*^−/−^ mutants and WT brains exhibiting largely similar spatial–topo tradeoffs across baseline and PTZ conditions. (A) A network was generated and a range of parameters was tested with (*η*, cost) ranging from -1.5 to 0 and (*γ*, reward) from 0 to 1 based on the probability produced in Eq. 1. Simulated networks were created, and the simulation was stopped when the network’s connections reach 10% of the total possible connections (30 edges shown here for illustration). (B) Each set of *η* and *γ* was repeated 300 times with random initial conditions. Each simulation was evaluated against the functional network derived from empirical calcium imaging data (top 10% correlation of each fish at each PTZ time) using KS energy similarity (Eq. 2, Figure S5). Higher similarity is indicated in yellow on the parameter landscape. (C) Parameter landscapes (a.k.a fingerprints) for each genotype at the pre-PTZ baseline show the best-fit values of *η* and *γ* that yield the lowest deviation from our empirical data. Red asterisks indicate the global minimum (best fit) and red dashed lines indicate the areas that are within top 10% of the best matches. (D) Comparison of the best-fit parameters (red asterisks in B) from the whole-brain coarse-grain modeling for each fish through the experiment, with each dot representing a single fish, while the line and shaded area indicating mean and SEM. Blue indicates the WT group, while red represents the *scn1lab*^−/−^ group. Effect sizes for genotype differences were quantified using Cohen’s *d* and *η*^2^, providing parameter-level estimates of divergence. (E) Relative amplitudes of *η* and *γ* throughout the experiment, where blue circles indicate that the WT mean is greater and a red circle indicates that the *scn1lab*^−/−^ mean is greater. The black ring denotes statistically significant results where p *<* 0.05, as determined by rANOVA.

We next selected 16 regions implicated in our graph theory analysis, and conducted detailed simulations for these regions. For each region, we performed the same GNM analysis, but at single-cell resolution, using *η* values ranging from -1.5 to 0 and *γ* values from 0 to 1. We ran 100 simulations for each pair of values, thereby identifying the optimal *η* and *γ* values for each genotype at each time point in the experiment. In the pallium (Figure 7A), and especially the habenula (Figure 7B), the optimal | *η* | and | *γ* | were consistently and significantly higher in WT than in *scn1lab*^−/−^. This could suggest that connections in WT are more structured, whereas *scn1lab*^−/−^ counterparts exhibit a higher degree of randomness in their network formation. The EG shows patterns where the optimal | *η* |, and to a lesser degree |*γ* |, values are higher in WT at pre-PTZ but shift to being higher in *scn1lab*^−/−^ following PTZ application (Figure 7B).

**Figure 7.**
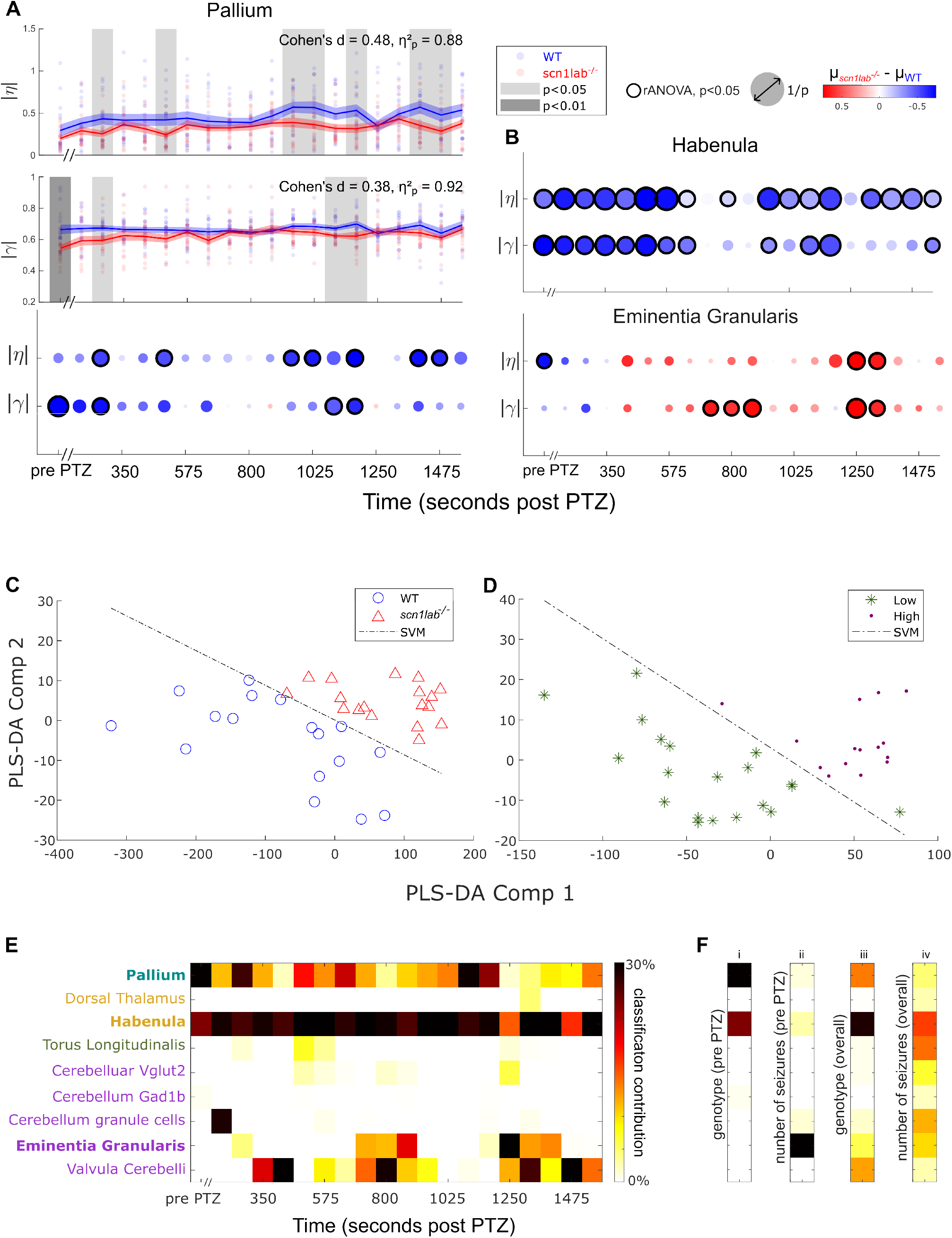
PLS-DA classifications using GNM fingerprints. Local single-cell modeling reveals genotype-specific parameter differences in select brain regions, and PLS-DA using subregional fingerprints enables accurate classification of both genotype and seizure severity, highlighting predictive features embedded in pre-PTZ network organization. (A) Best-Fit parameter comparison for local circuitry simulations of pallium at cellular resolution. Top panels show relative amplitudes of *η* and *γ* across the experiment; each dot represents an individual fish, with lines and shading indicating group mean and SEM. WT and *scn1lab*^−/−^ genotypes are shown in blue and red, respectively. Gray shading marks statistically significant differences (p *<* 0.05, repeated-measures ANOVA). Effect sizes were estimated using Cohen’s *d* and *η*^2^. Bottom panel summarizes genotype-level contrasts. (B) Genotype comparisons revealed significant differences in best-fit parameters within both the habenula and eminentia granularis (EG). For both (A) and (B), blue circles indicate a higher WT mean, while red circles indicate a higher *scn1lab*^−/−^ mean. A black ring marks statistically significant differences (p *<* 0.05), determined by rANOVA. (C) Using fingerprints from each simulation (Figure 6C) as features, PLS-DA effectively separates genotypes based on the 16 pre-PTZ subregion models. Blue circles represent WT animals, and red triangles represent *scn1lab*^−/−^ animals. The Support vector machine (SVM) line is drawn to demonstrate the genotype clusters can be linearly separated. (D) PLS-DA can also separate high and low numbers of seizures based on pre-PTZ subregions’ fingerprints. Green asterisks represents the low seizure count group (*<* 5.5 incidents), and magenta represents the high seizure count group (≥ 5.5 incidents). The linear SVM is also drawn to demonstrate the clustering separation. (E) The top 10% of contributions from each PTZ time point are identified to enhance genotype classification accuracy. Darker colors indicate a higher regional contribution to correct genotype classification. (F) The top 10% overall contributions from each subregion highlight accurate classification for (i) genotype from pre-PTZ data (as in C), (ii) number of seizures from pre-PTZ data (as in D), (iii) genotype from overall data (as in Figure S7), and (iv) number of seizures from overall data (as in Figure S7).

To evaluate the reliability of the generative model, we applied principal component analysis (PCA) to assess the stability of parameter estimates across all fish. Figure S6 shows that the individual effects of *η* and *γ* vary across runs, with neither parameter alone providing consistent clustering. However, when analyzed jointly (Figure S6), their combined variance uncovers structured separation among brain subregions, suggesting meaningful parameter recovery. In this plot, each mark represents an individual fish (n = 35), consistently grouped by subregion, emphasizing the reliability and stability of the GNM model at the brain region level. This separation correlates with anatomical proximity (Figure S6), reinforcing the biological plausibility of the fitted parameters. Notably, we identify reduced regulations on the wiring organization for *scn1lab*^−/−^ brains, reflected by more diffuse and overlapping PCA distributions, while WT brains show clearer spatial differentiation. This PCA analysis highlights that the genotypes exhibit distinct wiring principles, with some regions displaying more pronounced differences than others.

Our PCA analysis of the parameters *η* and *γ* uncovered distinct regional profiles between WT and *scn1lab*^−/−^ animals (Figure S6), prompting further classification using partial least squares-discriminant analysis (PLS-DA). This approach uses regional GNM fingerprints to optimize covariance with biologically relevant labels including genotype, seizure frequency, and network synchronization. By assessing all 16 regional models across 20 temporal windows, we identified the subregions most predictive of these phenotypes (Figure 7C–F), revealing them as key contributors to seizure risk.

Figure 7C shows two distinct clusters, demonstrating a clear separation between WT and *scn1lab*^−/−^ genotypes based solely on the fingerprints obtained from the pre-PTZ models. Similarly, Figure 7D reveals two well-defined clusters, indicating a strong distinction between high and low seizure count phenotypes, also derived exclusively from pre-PTZ models. The classification accuracy of the model ranges from 70–80%, as validated by a 10-fold cross-validation with 15 repeats (Figure S7). This demonstrates the effectiveness of GNM, applied to cellular-resolution functional data, to uncover baseline network properties inherent to *scn1lab*^−/−^ larvae and predictive of seizure susceptibility.

Because the PLS-DA analysis was performed on fingerprints for numerous brain regions implicated in seizures, we can identify which regions made the strongest contributions to the GNM’s discriminatory power (Figures 7E and 7F). This analysis implicates the pallium and habenula as regions whose pre-PTZ properties differ from WT in a way that correlates strongly with the animals’ genotypes (Figure 7Fi). In contrast, baseline network features in the cerebellar EG carried the greatest predictive power for post-PTZ seizure counts (Figure 7Fii, S7). For this analysis, each animal was labeled as either low- or high-seizure groups based on its seizure count. Expanding the classification to include multiple seizure count levels confirmed the EG as the dominant contributor, with additional predictive input from the habenula and cerebellar granule cells (Figure S7).

When taking information from across the entire experiment (pre- and post-PTZ) into account, the pallium and habenula are consistently strong indicators of genotype, with meaningful input from the cerebellum’s valvula cerebelli and, to a lesser degree, other cerebellar regions (Figures 7E and 7Fiii). Across the whole experiment, network parameters from most of the 16 sub-regions, especially from the habenula, can be used to predict the seizure counts for each animal (Figure 7Fiv). Additionally, two regions within the cerebellum, the *gad1b*- and *vglut2*-positive clusters, emerge as key regulators of brain-wide network synchrony, such that the overall synchrony can be predicted using parameters from these regions alone (Figure S7). Overall, we identify the brain regions and network properties that predispose *scn1lab*^−/−^ animals to seizures, using approaches that were reliant on cellular-resolution input data.

## DISCUSSION

In this study we have investigated the genotype-driven changes in functional neural network structure by comparing *scn1lab*^−/−^ mutant and WT zebrafish larvae under PTZ-induced seizure conditions. Our analysis has revealed genotype-dependent morphological differences reflecting altered neuronal density. These structural changes were accompanied by widespread disruptions in whole-brain network synchrony, as evidenced by altered correlation patterns across hemispheres and within local regions. To pinpoint the functional drivers of these network shifts, we applied graph theoretic analyses and constructed GNMs using data at single-cell resolution. This multi-scale modeling strategy permitted accurate genotyping and prediction of seizure susceptibility in the absence of observable phenotypes or changes in broad activity-based metrics. These results suggest that the observed seizure susceptibility rests on changes in wiring rules, at the scale of whole networks but the resolution of neuron-to-neuron functional connectivity, that have traditionally been impossible to observe or assess. Our findings highlight the critical importance of cellular-resolution data, as these discrete spatial and functional features are lost when cellular-resolution data are aggregated or averaged.

### *scn1lab*^−/−^ zebrafish have altered brain morphology, functional wiring, and seizure susceptibility

We first examined morphological and baseline functional alterations in *scn1lab*^−/−^ mutants to assess their contribution to seizure susceptibility (Figure 3B-D). Mutant larvae exhibited reduced telencephalon and diencephalon volumes alongside rhombencephalic enlargement. ROI counts, inferred from ROI detection via Suite2p^29^, mirrored these anatomical changes: WT larvae showed higher ROI density in forebrain regions, whereas *scn1lab*^−/−^ mutants displayed enrichment in the hindbrain. There are several potential functional impacts of these morphological changes. A reduced density in the pallium and habenula could be a sign of altered monoaminergic signaling that is important to seizure suppression^49^. More broadly, reductions in the telencephalon and diencephalon could impair GABAergic inhibitory drive and alter sensorimotor gating, thus lowering seizure thresholds^50–52^. Although ROI counts are commonly interpreted as a proxy for neuron number, it is important to consider that Suite2p relies on calcium-dependent fluorescence for segmentation, potentially biasing detection toward more active neurons^29^. As such, reduced ROI counts in regions like the habenula of *scn1lab*^−/−^ larvae may reflect diminished activity rather than true cellular loss, while elevated counts in the hindbrain may instead indicate hyper-activity rather than increased cell density.

To probe the network-level consequences of these altered cell numbers, we characterized functional connectivity in both WT and *scn1lab*^−/−^ mutants under baseline and ictogenic conditions (Figure 3E, F). PTZ exposure increased whole-brain synchrony in *scn1lab*^−/−^ mutants, reflected in elevated correlation and reduced skewness (Figure S1C), consistent with prior regional findings^37^, but now contextualized at the network scale. These results allowed us to identify where and how the network changes in *scn1lab*^−/−^ mutants. Notably, we observed increased contralateral functional connectivity in *scn1lab*^−/−^ mutants, indicative of strengthened bilateral communication, presumably via commissural pathways (Figures 4 and S3A). This enhancement was evident even prior to PTZ exposure, particularly within the mesencephalon and rhombencephalon, counteracting the typical distance-dependent decline in correlation strength^38^. These findings suggest that excitation in one hemisphere could more readily propagate across the midline^53^, a phenomenon that intensifies with PTZ and may underlie seizure initiation and contralateral spread^54^. A similar mechanism is also observed in rodent models of temporal lobe epilepsy^55^, where commissural granule cell axons, normally restricted to unilateral projections, exhibit rapid contralateral growth, providing a structural basis for enhanced interhemispheric connectivity. This also mirrors secondary generalization of seizures in humans, where focal seizures recruit bilateral networks, manifesting as contralateral spread^56^.

Consistent with these interpretations of the altered anatomy and functional connectivity, we observed a significantly greater PTZ sensitivity and seizure risks in the *scn1lab*^−/−^ animals. Although neither genotype had spontaneous seizures, *scn1lab*^−/−^ larvae had more induced seizures and shorter latencies than WT (Figure 1C-E). Given PTZ’s broad suppression of inhibitory signaling^54^, these findings suggest that *scn1lab*^−/−^ brains have altered baseline network configurations that fundamentally increase seizure susceptibility and enhance synchrony at the whole network scale.

### PTZ impacts network structure and function differently in *scn1lab*^−/−^ and WT larvae

We assessed network-scale graph metrics (clustering coefficient, modularity, efficiency, and assortativity) that are critical for moving beyond descriptive correlation measures to identify the architectural principles underlying the seizure-prone phenotype (Figure 5A). At the pre-PTZ stage, *scn1lab*^−/−^ brains have higher assortativity, i.e., a greater tendency for nodes to connect with others of similar degree. This architecture, characterized by highly interconnected, resilient core networks, could facilitate rapid, synchronous propagation of abnormal activity, which is a hallmark of epileptic seizures^57^, and aligns with previous reports of pathophysiological conditions^41^. This difference in baseline network organization provides an explanation for the enhanced seizure susceptibility observed in *scn1lab*^−/−^, as the interconnected nature of these nodes could enable the rapid spread of epileptiform activity^58^.

We found several PTZ-induced changes, including higher means in WT for assortativity, efficiency, and modularity (Figure 5A). This constellation of features reflects a modular yet locally cohesive structure, characterized by tightly interconnected clusters. Notably, this PTZ-induced configuration in WT mirrors the baseline network topology observed in *scn1lab*^−/−^ mutants, which exhibit elevated assortativity in the absence of PTZ. Assortative networks, defined by preferential connectivity among nodes of similar degree, are known to facilitate the propagation of abnormal activity through hub-like structures (high modularity)^44^. The transition from non-assortative to assortative mixing under PTZ exposure compromises network resilience, rendering it more susceptible to structural perturbations such as vertex removal, and amplifying vulnerability to seizure-like disruptions^59^.

Elevated edge length values following PTZ administration reflect increased seizure events in *scn1lab*^−/−^ larvae (Figure 5A, B). These prolonged edge lengths are not only indicative of seizure episodes, but also point to a systemic reorganization of functional network architecture relative to baseline. We observed that seizure-associated hyper-synchrony facilitates aberrant long-range signal propagation across the zebrafish brain, resulting in a measurable decrease in network efficiency. These data suggest that the *scn1lab*^−/−^ mutation confers a latent topological vulnerability, where the transition to an ictal state is defined by a large-scale change of functional connectivity. Crucially, the use of cellular-resolution data reveals that the baseline *scn1lab*^−/−^ brain possesses an underlying structural resemblance to the PTZ-perturbed state, a signature of susceptibility that is lost in coarser data modalities.

These findings align with human studies of epileptic networks where there are significant changes to networks structures compared to controls. For example, Liao et al. (2010) reported that regions with reduced connectivity in Mesial Temporal Lobe Epilepsy (mTLE) patients were predominantly components of the default-mode network, and exhibited significantly lower normalized edge lengths^42^. Similarly, Pedersen et al. (2015) found that individuals with focal epilepsy displayed elevated clustering coefficients, local efficiency, and modularity relative to healthy controls^60^. Together, these graph theory metrics point to a shared vulnerability of the neural network to hyper-synchronization and disrupted information flow. This altered topology, observed in *scn1lab*^−/−^ mutants, reflects a broader principle of seizure susceptibility where tightly clustered yet poorly integrated networks are prone to pathological hyper-synchronization and inefficient signal delivery. Crucially, network vulnerabilities in *scn1lab*^−/−^ mutants are not randomly distributed but are systematically anchored to node degree alterations. This impairment in functional connectivity may represent a core network deficit underlying the emergent seizure phenotype.

Building on previous results and studies, our use of single-cell resolution imaging enabled precise localization of genotype-dependent network alterations across distinct brain subregions. To dissect these effects, we applied a node-wise metric, degree, following PTZ exposure (Figure 5C), allowing us to quantify region-specific changes in functional connectivity. Notably, a subset of brain regions, including the forebrain and midbrain, exhibited distinct degree changes compared to the hindbrain, reflecting region-specific vulnerability to PTZ-induced network disruptions. These findings suggest differential susceptibility to PTZ-induced hyper-excitability across anatomical domains, implicating specific subregions in the emergence of seizure phenotypes. Importantly, these connectivity-based alterations align with morphological and ROI count differences observed in mutants (Figure 3B-D), reinforcing the notion that genotype-driven structural changes contribute to defects in the network structure-function relationship that increase seizure risk.

### GNM reveals the brain regions and network architecture responsible for seizure risk

GNM offers a principled approach for understanding the evolution of the architecture of functional brain networks by simulating their formation using biologically plausible rules. Rather than reconstructing time-resolved activity, which is often computationally demanding, GNM infers network organization based on structural connectivity reflecting the assumption that functional interactions emerge from underlying pathways or structures^61–63^. While GNM does not create identical networks, it preserves key functional properties measurable by graph theory. In this framework, structure–function relationships are assessed at the network level, where the topology of structural connections shapes the emergence of functional connectivity patterns, quantified using graph theory metrics.

GNM is empirically constrained and involves minimal assumptions, yet incorporates physiological explanations, making it especially suited for exploring large-scale differences in functional brain networks such as those observed in our dataset. A key strength of GNM is its reliance on graph theory comparisons between simulated and empirical networks, which helps preserve core properties like efficiency and degree without requiring the simulation of full time series. This mitigates bias toward any single measurement and supports a generalizable analysis framework^45^. While simulated networks may not replicate empirical connectivity patterns exactly, their topological similarity suggests convergence on shared organizational principles. The trade-off between biological plausibility and computational efficiency enables meaningful insights into the mechanisms shaping functional network organization across individuals or groups.

In using GNM to describe network dynamics in our model of epilepsy, we initially employed a coarse-grained approach to examine whole-brain network formation. While this method yielded an optimized simulated network, it failed to reveal significant differences in their best-fit parameters *η* and *γ* between genotypes at the whole-brain level (Figure 6D, E). This absence of pronounced phenotypic differences in coarse-grained data, reminiscent of challenges encountered in fMRI studies, highlights the complexity and importance of local networks and the limitations of coarse-grained approaches in capturing subtle, yet potentially crucial, alterations in brain functional connectivity.

To overcome these limitations, we turned to cellular-resolution data, focusing on specific brain subregions. To ensure an unbiased selection, 16 subregions chosen for GNM modeling were solely based on findings from the first part of this study. This granular approach revealed genotype-dependent alterations in the functional architecture of neural networks between WT and *scn1lab*^−/−^ mutant animals. Particularly in the pallium (Figure 7A), we observed higher *η* and *γ* values in WT compared to *scn1lab*^−/−^, indicating stronger governing parameters for both cost (Euclidean distance) and reward (matching index) in WT networks. These elevated values in WT suggest a more selective and spatially constrained connectivity regime, favoring proximity and topological similarity. In contrast, *scn1lab*^−/−^ networks displayed reduced parameter strength, indicative of a more stochastic and less organized wiring strategy. This discovery aligns with previous research linking wiring costs to polygenic risk for schizophrenia^64^ and Alzheimer’s disease^43^, suggesting that the network bases for a diversity of neurological conditions can be detected as alterations in GNM parameters.

To validate the model and examine the effects of its parameters, we applied PCA to the bestfit parameters to explore internal relationships between subregions and genotypes (Figure S6). Proximal regions displayed similar wiring parameters, suggesting consistent rules for connection formation. While individual *η* and *γ* parameters showed subtle genotypic differences, their interaction was crucial in distinguishing the formation rules from *scn1lab*^−/−^ and WT groups. Notably, *scn1lab*^−/−^ brains displayed less differentiated wiring clusters among different regions, particularly in the rhombencephalon. These findings suggest that the fundamental connectivity principles in *scn1lab*^−/−^ brains are altered, potentially reducing functional specialization across brain regions and increasing seizure susceptibility ^65,66^. Furthermore, these results demonstrate that PCA-based validation provides insight into the robustness and fidelity of model fitting, supporting its use in connectomic inference.

Our analysis further revealed a compelling mathematical signature of seizure risk, with specific brain regions demonstrating remarkably robust predictive power for both genotype and seizure risk phenotype, based solely on baseline (pre-PTZ) data (Figure 7E, F). Key subregions such as the habenula, pallium, EG, and valvula cerebelli emerged as critical contributors to the accurate classification of genotypes and prediction of seizure counts. Notably, the EG stood out as the primary contributor to seizure count predictions, suggesting a direct link between its altered connectivity patterns and ictogenesis in *scn1lab*^−/−^ mutants. The valvula cerebelli’s importance in distinguishing genotypes during seizure activity, despite lacking clear morphological distortions or changes in GNM parameters, implies that altered micro-connectivity produces the mutants’ susceptibility to seizures and highlights the benefits of a cell-resolution approach.

Using the same approach, we also examined which subregions drive changes in network correlation. By categorizing the mean network correlation values into five bins, we identified key subregions influencing the fluctuations in network synchronization (Figure S7). Neurons found in the *gad1b*- and *vglut2*-expressing regions of the cerebellum were the strongest contributors to accurate classification of correlation values. GNM-derived best-fit parameters revealed significantly elevated WT values for both *η* and *γ* at baseline in the *gad1b*-expressing region, which modulates GABA co-transmission^67^. These findings suggest that the mutation may disrupt GABAergic wiring principles, rendering *scn1lab*^−/−^ networks more susceptible to PTZ-induced perturbation. In contrast, prior to PTZ exposure, no genotype-dependent differences in *η* or *γ* were observed in the *vglut2*-expressing region, which regulates excitatory glutamate signaling^67^. Following PTZ administration, however, *γ* values in the *scn1lab*^−/−^ networks increased markedly in this region, particularly during seizure-dense epochs. This effect is consistent with PTZ-enhanced glutamate release^17^, which may elevate homophily (cellular matching index) and promote hyper-synchronization. As a caveat, it is important to note that, while these regions within the cerebellum express *gad1b* and *vglut2*, they are occupied by heterogeneous neurons, not all of which express the identifying marker. The absence of *η* changes indicates that inter-cellular distance constraints (*D*_*ij*_) remained stable across genotypes.

These results suggest that the cerebellum’s value for predicting seizure susceptibility rests in its regulating the brain’s excitatory/inhibitory (E/I) balance (Figure 7F). Specifically, under normal conditions, WT inhibitory neurons exhibit stronger regulatory control compared to their counterparts in *scn1lab*^−/−^. We propose that, following the application of PTZ, a noncompetitive GABA_*A*_ receptor antagonist^68^, GABA neurons in *scn1lab*^−/−^, which already exhibit reduced activity, rapidly fall below the inhibitory drive threshold, triggering seizures. Prolonged exposure to PTZ, on the other hand, effectively suppresses GABA signaling in WT, ultimately initiating seizures in WT as well^54^. These interpretations align with the convergence of GNM parameter values observed later in the experiment, when both genotypes experience seizure activity. Increased *γ* values during seizure-intensive periods in *vglut2*-expressing regions suggest corresponding dynamics, with nodes that are typically unconnected in WT becoming readily interconnected in the *scn1lab*^−/−^ mutants, or only after a persistent exposure to PTZ in WT. As noted above, because our registration is based on spatial location rather than molecular identity, not all neurons necessarily express the neurotransmitter typical of their region, and our data do not allow neurotransmitter subtyping at cellular resolution. While the cerebellum may show spatial segregation of excitatory and inhibitory populations, such organization may exist in other brain regions, but without observable spatial separation. Future experiments that employ antibody labeling or transgenic methods to definitively identify the neurotransmitter types of individual neurons will help to corroborate or elaborate on the seizure network’s underlying biological mechanism.

### Methodological perspective and future directions

Previous studies of epilepsy have emphasized how *SCN1A* mutations alter neuronal excitability, leading to hyper-excitable networks and spontaneous seizures^69,70^. Functional imaging modalities such as fMRI and EEG have revealed widespread alterations in brain activity^69,71^, suggesting that seizure networks extend beyond focal excitability changes. Electrophysiological recordings have further characterized seizure dynamics^34^, though they lack the multi-scale resolution to reveal the broader connectivity patterns critical to ictogenesis.

Zebrafish carrying *scn1lab*^−/−^ mutations offer complementary insights into seizure mechanisms by bridging gaps left by conventional imaging and electrophysiological approaches. For instance, Yaksi et al. (2021) have demonstrated that *scn1lab*^−/−^ mutants exhibit impaired inhibitory control and GABAergic dysfunction^20^. The zebrafish’s optical accessibility and genetic tractability enable in-depth exploration of seizure dynamics using whole-brain calcium imaging, bridging whole-network activity tracking with single-cell resolution^20^. These approaches support a systems-level view of epilepsy, implicating excitatory/inhibitory imbalance^72^, altered network topology^73^, and disrupted global connectivity^74^.

Rodent and human studies further reinforce this perspective. Wong et al. ^75^ (2016) showed that *Scn1a*^+/−^ and *Scn1a*^*RH/*+^ mice are highly sensitive to PTZ-induced seizures, with Huperzine A conferring protection via muscarinic and GABA_*A*_ receptor modulation^75^. Similarly, Das et al. ^76^ (2021) reported interneuron dysfunction in *Scn1a*^*KT/*+^ mice, mirroring the hyper-excitable states observed in PTZ-challenged zebrafish^76^. In clinical contexts, *SCN1A* mutations are linked to febrile seizures and pharmacoresistant epilepsy^77^, primarily due to Na_V_1.1 channel dysfunction that disrupts inhibitory interneuron activity and skews excitability balance^77^.

Our study builds on these findings by integrating whole-brain calcium imaging with a multiscale network analysis including GNM to characterize seizure-prone architectures in *scn1lab*^−/−^ mutants. We observed increased synchronization, disrupted power-law decay, and enhanced mid-hindbrain connectivity—features indicative of pathological network reorganization^19,45^. GNM confirmed that these changes arise from spatial and functional constraints on connectivity, producing networks with reduced robustness and heightened susceptibility to pharmacological challenge. A key strength of our approach lies in the use of cellular-resolution imaging to uncover network-level differences that remain obscured in coarse-grained analyses. While baseline activity appeared comparable across genotypes, the introduction of PTZ revealed striking divergence in single-cell responses, particularly within distinct brain regions. These region-specific perturbation profiles were not detectable at the population level, highlighting the importance of spatial granularity. Moreover, by probing network stability through PTZ-induced dynamics, we were able to infer genotype specific vulnerabilities, such as those in *scn1lab*^−/−^ mutants, which would be inaccessible without both perturbation and cellular resolution data.

By identifying network-scale alterations in wiring rules that underlie seizure susceptibility, this approach will enable comparative analyses across genetic epilepsy models, including rodent *Scn1a* mutants, and may reveal conserved principles of circuit instability^20^. Future studies incorporating patient-derived alleles and broader perturbation paradigms could refine genotype–phenotype associations and inform precision strategies for intervention.

Epilepsy is widely recognized as a heterogeneous disorder at the whole-brain level, yet most existing models rely on mean-field or population-level dynamics and connectivity^78^ that obscure individual variability in circuit function^20^. Our approach addresses this gap by enabling multiscale analysis of seizure-related network dynamics at cellular resolution. By integrating functional connectivity with a computational modeling approach, we can now quantify genotype-specific differences within individual network organization and perturbation responses, features that are currently not possible in human studies. This zebrafish model, with its optical transparency and genetic tractability, offers a uniquely scalable platform for resolving these differences across individuals and genotypes. In particular, the network’s response to PTZ reveals latent circuit vulnerabilities that predict seizure susceptibility, providing a mechanistic link between genotype and phenotype. This framework moves beyond coarse-grained mean-field models to capture the diversity of epileptic network topology and states, offering new opportunities for precision, individually specific modeling and therapeutic targeting.

## Supporting information

Video 1. Behavioral manifestations observed in zebrafish during seizure development.

Video 2. An animated video showing comparison between SyN and ROI density over different planes

Video 3. An animated video illustrates the temporal dynamics of correlation strength across ipsilateral and contralateral connections.

## ONLINE METHODS

### Experimental setup and data acquisition

#### Experimental setup

All experimental procedures, including the housing and breeding of adult zebrafish as depicted in Figure S1, were conducted in strict compliance with the guidelines approved by the University of Queensland Animal Ethics Committee (Approval No. 2021/AE001047). Adult zebrafish were accommodated at a density ranging from 10 to 15 fish per Liter and were subjected to a 14/10-hour light/dark cycle at a consistent temperature of 28.5°C.

The breeding involved an incross of nacre *scn1lab*Δ*44* heterozygous specimens carrying the elavl3:H2B-GCaMP6s transgene, resulting in a Mendelian distribution of homozygous (*scn1lab*^−/−^) heterozygous (*scn1lab*^−/+^), and Wild-Type (*scn1lab*^+/+^) embryos in a ratio of 1:2:1. Each clutch comprised 100 fertilized eggs, which were incubated at 28.5°C in Petri dishes filled with E3 medium. The E3 medium consisted of distilled water supplemented with 10% Hanks solution, containing the following concentrations:

- NaCl: 137 mM
- KCl: 5.4 mM
- Na_2_HPO_4_: 0.25 mM
- KH_2_PO_4_: 0.44 mM
- CaCl_2_: 1.3 mM
- MgSO_4_: 1.0 mM
- NaHCO_3_: 4.2 mM

The medium was maintained at a pH of 7.2. At four days post-fertilization (4dpf), larvae were assessed for GCaMP6s expression. Those exhibiting positive expression were selected for calcium imaging experiments conducted at six days post-fertilization (6dpf). Following the experiments, the larvae were euthanized and fixed in paraformaldehyde (PFA) overnight. The samples were then washed with phosphate-buffered saline (PBS) and stored at 4°C. The heads, still embedded in agarose and PBS, were preserved for future research, while the tails were digested with proteinase K. The DNA extracted from the tails was subsequently amplified through polymerase chain reaction (PCR) for genotyping. Post-experimental genotyping of the larvae was performed using gel electrophoresis to identify the *scn1lab*^−/−^ allele at 372 base pairs (bp) and the WT allele at 416 bp.

Out of the initial 68 larvae subjected to the 5 mM PTZ paradigm, the final dataset was refined to exclude 21 heterozygotes, 3 specimens that expired during the experiment, and 9 due to suboptimal data quality, such as inadequate focus or excessive tilting beyond the capacity for warp correction. Consequently, the analyzed dataset comprised 18 homozygous *scn1lab*^−/−^ larvae and 17 Wild-Type counterparts.

#### SPIM and setup

The schematic representation of the calcium imaging experiments provides a comprehensive overview of the setup (Figure S1,). The side view illustration of the imaging chamber reveals the precise alignment of the orthogonal imaging objective, which is meticulously focused on the fish’s brain. Additionally, a separate camera is positioned beneath the chamber, dedicated to the precise tracking of tail movements. From the top perspective, the imaging chamber is depicted with two distinct light sheets strategically positioned for optimal imaging: one oriented anteriorly towards the front of the fish and the other laterally from the side.

Larvae were immobilized using 2% low melting point agarose and positioned vertically in a 7mL imaging chamber^32^. The agarose surrounding the tail was carefully removed, securing the head while allowing tail mobility (by removing segments of agarose perpendicular to the tail caudal to the swim bladder, and the chamber was filled with E3 media), as illustrated in Figure 1A. A minimum acclimatization period of one hour was allotted to ensure the larvae were settled into the agarose, minimizing Z-axis drift during imaging.

Tail movement (Figure 1B) was concurrently imaged using infrared illumination and a FLIR Blackfly USB camera situated beneath the chamber operating at 200 frames per second (fps), and synchronized with the calcium imaging through a timed infrared LED flash. The tail was tracked using a spline-fitting algorithm^79^.

Calcium imaging recordings were acquired from 17 wild-type (WT) and 18 *scn1lab*^−/−^ mutant zebrafish, encompassing a 5-minute baseline period prior to pentylenetetrazol (PTZ) administration and a 30-minute post-PTZ observation window. The experimental timeline was segmented into three distinct phases: pre-PTZ (baseline spontaneous activity), early-PTZ (0–15 minutes post-administration, characterized by moderate seizure frequency), and late-PTZ (15–30 minutes post-administration, marked by elevated seizure activity). No behavioral or physiological recovery was observed during the post-PTZ interval.

Calcium imaging data were acquired at 2 Hz and processed to correct for motion artifacts, delineate regions of interest (ROIs), and compute normalized fluorescence changes (Δ*F/F*_0_). On average, ∼20,000 active ROIs were identified per specimen across 25 imaging planes. ROI centroids were registered to the Zbrain atlas to enable standardized anatomical referencing and facilitate cross-specimen comparisons.

Calcium imaging data (Figure S1) were captured using a custom-built one-photon selective plane illumination microscope^80^. Briefly, dual 488 nm lasers concurrently illuminated the brain in a narrow horizontal plane, facilitating the visualization of neuron activity via the genetically encoded calcium indicators (GECIs), *elavl3*:H2B-GCaMP6s, through an orthogonal objective lens. Images were captured at a resolution of 640 × 540 pixels, binned four times, and saved in 16-bit TIFF format. The light sheet traversed the Z-axis in 10*µm* increments at 2 Hz, generating 25 horizontal planes per specimen to compile a comprehensive volumetric representation of the entire brain. This method provides a quantitative measure of calcium concentration changes within the cells at the cellular resolution, reflecting neuronal activity.

#### Calculation of PTZ stock concentration

Imaging chambers were filled with 7 mL of E3 medium. To achieve a final pentylenetetrazol (PTZ) concentration of 5 mM, 1 mL of E3 was removed and replaced with 1 mL of PTZ stock solution, following the standardized experimental protocol. For preparation of the working solution, a 35 mM PTZ stock was generated, from which 1 mL was added to 6 mL of E3 medium to yield 7 mL of 5 mM PTZ. This concentration was selected to reliably induce seizure-like activity while maintaining viability throughout the imaging session.

#### Extraction and analysis of fluorescence traces

ROIs, each representing a single neuron, were identified and spatially registered in three dimensions using a custom Python-based pipeline. This workflow incorporated Suite2p (v0.14.3) for ROI extraction and Advanced Normalization Tools (ANTs, v2.4.4) for image alignment, executed on the Spartan high-performance computing (HPC) cluster at The University of Melbourne^81^. Fluorescence traces (Δ*F/F*_0_) were computed for each ROI to quantify neuronal activity (Figure S1). Suite2p identified candidate signal sources by detecting pixels with high multidimensional correlation within localized spatial neighborhoods^29^. ROIs were classified using a custom-trained algorithm developed in-house, and fluorescence signals were computed via weighted averaging of constituent pixels, with correction for neuropil contamination. Motion correction was performed automatically during Suite2p registration^33^. Suite2p configuration parameters are listed in Table 1. ROI outputs and corresponding traces were subsequently analyzed in MATLAB^82^.

**Table 1:** Suite2p Configuration Parameters.

| Parameter | Value |
| --- | --- |
| nplanes | 25 |
| tau | 1.4 |
| fps | 100 |
| sampling frequency | 2 |
| save_mat | 1 |
| batch_size | 2000 |
| 1Preg | true |
| pre_smooth | 2 |
| spatial_taper | 5 |
| diameter | [3, 4] |
| threshold_scaling | 0.5 |
| max_iterations | 100 |
| inner_neuropil_radius | 1 |
| min_neuropil_pixels | 1 |
| sparse_mode | false |
| spikedetect | false |
| smooth_sigma | 4 |
| high_pass | 30 |
| combined | false |
| classifier_path | ~/hpc_pipeline/classifier.npy |

**Table 2:** Graph theory metrics with mathematical meanings and biological interpretations.

| Metric | Mathematical Meaning | Biological Interpretation |
| --- | --- | --- |
| **Degree** | Number of direct connections a node has; reflects local connectivity. | Indicates how many functional connections a neuron maintains; high degree may suggest hub-like behavior or hyper-connectivity. |
| **Path Length** | Average shortest distance between nodes; relates to network navigability. | Reflects how efficiently signals can propagate across the brain; shorter paths imply faster communication. |
| **Topological Overlap** | Measures shared neighbors between node pairs; captures local redundancy. | Suggests coordinated or redundant connectivity among neurons; increased values may indicate robust or synchronized network modules. |
| **Betweenness Centrality** | Fraction of shortest paths passing through a node; identifies control points. | Highlights neurons that act as communication bottlenecks or integrators across brain regions. |
| **Assortativity** | Correlation between degrees of connected nodes; measures preference for similar connectivity. | Indicates whether highly connected neurons preferentially link to other hubs; may reflect developmental or pathological wiring biases. |
| **Clustering Coefficient** | Likelihood that a node's neighbors are also connected; quantifies local cliquishness. | Captures local modularity or segregation; high values suggest tightly-knit functional communities. |
| **Efficiency** | Inverse of path length; measures global and local information transfer. | Reflects how effectively the brain network supports rapid and integrated communication. |
| **Modularity** | Degree to which the network can be partitioned into distinct communities. | Reveals functional specialization across brain regions; high modularity may indicate compartmentalized processing. |

To enable anatomical standardization, ROI centroids were registered to Zbrain atlas using Advanced Normalization Tools (ANTs)^30,35^ following previously published method^33^. Registration employed predefined similarity metrics to normalize spatial coordinates and correct for interindividual variation in orientation. A high-resolution template stack (1 *µm* z-spacing), generated from motion-corrected, time-averaged Suite2p outputs, was aligned to the Zbrain atlas^32^. The resulting transformation matrices were applied to ROI centroids to ensure consistent spatial mapping across specimens.

To facilitate region-level analyses and reduce redundancy, a curated subset of 137 brain regions was derived from the original 295-region Zbrain atlas. This refinement merged overlapping subregions and eliminated unnecessary anatomical subdivisions. Warped ROI coordinates were assigned to one of the 137 standardized regions; ROIs located outside atlas boundaries following registration were excluded from downstream analysis.

### Mathematical methods

#### Jacobian analysis via SyN registration

To quantify volumetric deformation during atlas registration, we applied symmetric image normalization (SyN), a diffeomorphic mapping algorithm that optimizes cross-correlation using Euler–Lagrange equations within the ITK framework^36^. Originally developed for MRI-based morphometry, SyN enables precise spatial alignment by modeling expansion and contraction of anatomical regions. In our study, SyN was used to compare ROI coordinates pre- and post-registration with ANTs, highlighting regions requiring volumetric adjustment for accurate atlas alignment.

#### ROI density mapping

To assess genotype-specific differences in spatial ROI distribution, we projected all ROIs onto a standardized anatomical template and computed density within 6 ×6 ×6 *µm*^3^ cubes. Each brain volume was segmented into 148 × 90 × 17 cubes, avoiding double-counting across 10 *µm*-spaced planes. ROI density was defined as the mean number of ROIs per cube, and statistical comparisons between WT and *scn1lab*^−/−^ genotypes were performed using the Mann—Whitney U test.

Significant differences in ROI density were visualized per cube, with blue and red indicating higher density in WT and *scn1lab*^−/−^, respectively. Representative slices at 70 *µm* and 130 *µm* are shown in Figure 3B, with additional planes provided in Figure S2.

#### Seizure detection algorithm

Seizures are classically defined as “an excessive, hyper-synchronous discharge of neurons”^83^. To align with this definition while accommodating the spatiotemporal resolution of zebrafish calcium imaging, we design seizure detection using a fully automated, data-driven pipeline that integrates behavioral, neural, and computational metrics. Localized activity patterns (Figure S1) are designated as subthreshold ensembles, whereas widespread, synchronous activation across brain regions is classified as seizure events.

Seizure onset is identified by the concurrent emergence of four hallmark features: (1) behavioral hyper-activity, defined as tail angle excursions exceeding the 99.9^th^ percentile of the interquartile range (IQR); (2) elevated neural activity, defined as Δ*F/F*_0_ exceeding the 99^th^ percentile IQR across ROIs; (3) abrupt head movement, inferred from motion correction values exceeding the 99^th^ percentile IQR across imaging frames; and (4) network synchrony, quantified via skewness of the pairwise correlation distribution, with seizure-classified events exhibiting skewness *>* 0.8. The skewness threshold reflects the extent to which a majority of neurons exhibit hyper-synchronized activity, distinguishing network states where high correlations are concentrated among most ROIs, while only a minority remain weakly coupled.

Motion correction was performed using Suite2p, which estimates frame-to-frame lateral displacements (corrXY) to correct for movement artifacts across imaging planes. For each animal, motion correction traces were extracted from Suite2p’s Fall.mat files across all 25 imaging planes, and the XY displacement values were aggregated to generate a whole-brain motion profile. These values reflect the magnitude of lateral shifts applied during registration and serve as a proxy for physical movement or imaging instability. To enhance interpretability, the raw motion traces were baseline-adjusted using a custom differencing function, emphasizing transient motion bursts while suppressing slow drift.

Prior to PTZ administration, no seizure-classified events were detected in either genotype, and skewness values remained below threshold (Figure S1C). However, post-PTZ, both mean correlation and skewness metrics diverged significantly between genotypes, indicating altered network dynamics. Notably, *scn1lab*^−/−^ mutants exhibited elevated baseline neural activity relative to wild-type siblings, suggesting a heightened excitability state even in the absence of overt seizures.

#### Correlation strength comparison

To assess distance-dependent functional connectivity, Euclidean distances between all ROI pairs were binned in 100 *µm* intervals. Within each bin, mean correlation coefficients were computed and normalized to the 0-100 *µm* baseline, yielding correlation offsets relative to each animal’s pre-PTZ state. Genotype differences were evaluated across distance ranges using Mann-Whitney U tests (Figure 4).

For enhanced resolution, Figure S3 presents genotype-specific offsets at the region-to-hemisp and intra-regional levels using 30 *µm* bins, consistent with patterns in Figures 4C-D.

#### Edge density comparison

To complement correlation offset analysis, we quantified the number of highly correlated edges (top 10% of correlation values) across genotypes. Two network types were constructed: a wholebrain network from the global correlation matrix, and local subnetworks from subregion-specific matrices. In both cases, top correlations were binarized to generate adjacency matrices, and edge counts were compared per region.

The whole-brain approach binarizes the full matrix before extracting inter-regional edges, while the local approach computes and binarizes subregion-specific matrices directly. Wholebrain analysis highlights genotype-specific edge density across subregions (Figure S3), whereas local networks reveal intra-regional connectivity differences (Figure S3).

#### Graph theory

To assess genotype-specific differences in response to PTZ, we applied two complementary approaches: node-wise and network-wise graph metrics. Functional connectivity networks were constructed by thresholding Pearson’s correlation matrices at the top 10%, yielding binarized adjacency matrices. Metrics were computed using the Brain Connectivity Toolbox^40^.

Node-wise metrics, including degree, path length, topological overlap and betweenness centrality, were averaged across ROIs within each subregion and statistically compared between genotypes. Only subregions showing consistent differences across PTZ stages are shown (Figure S4).

Network-wise metrics focused on local connectivity within each subregion, independent of global topology. Metrics included assortativity, clustering coefficient, efficiency, and modularity, revealing genotype-specific alterations in local network organization (Figure S4).

#### Randomization and null models

To assess whether graph theory metrics reflect intrinsic network properties or arise from degree-related artifacts, Maslov–Sneppen (MS) randomization was applied to empirical adjacency matrices^84,85^. This degree-preserving edge-swapping procedure disrupts higher-order topology while maintaining local connectivity. For each zebrafish specimen, −100 MS-randomized surrogates were generated to construct null distributions for modularity, clustering, and other metrics. This approach enables statistical comparison between observed values and degree-controlled expectations, clarifying whether functional modules reflect biologically meaningful organization or trivial structural constraints.

To evaluate the temporal specificity of functional connectivity, phase randomization was applied to the fluorescence time series of each ROI. Each Δ*F/F*_0_ trace was transformed via fast Fourier transform (FFT), its phase spectrum randomized, and inverse-transformed to yield surrogate signals with preserved power spectra but disrupted temporal structure. These phase-randomized datasets were used to generate null models for pairwise correlation distributions (Figure 3Ei) and graph metrics (Figure S4), allowing us to distinguish genuine neuronal coactivation from artifacts introduced by spectral content or preprocessing. This method provides a rigorous control for assessing dynamic interactions in whole-brain zebrafish networks.

#### Generative network modeling (GNM)

Connectome generative models simulate brain network development by applying probabilistic wiring rules that balance spatial and topological constraints, offering a framework for studying cost-efficient architecture. When parameters are optimized to match empirical data, their variation across individuals reveals associations with aging^46^, cognition^45^, and neurodevelopmental adversity^46^.

GNM bypasses time-series fitting by directly matching simulated networks to empirical functional connectivity. Our implementation uses a single cost function (Euclidean distance between ROIs; Figure 4B) and a reward function, matching index, based on prior knowledge^45^. Connection probability is governed by:

- *P*_*ij*_: Probability of edge formation between nodes *i* and *j*
- *D*_*ij*_: Spatial cost (Euclidean distance)
- *K*_*ij*_: Topological similarity (matching index)

The scaling parameters *η* and *γ* modulate the influence of spatial cost and topological similarity, respectively. Positive values promote edge formation, negative values suppress it, and zero values exert no effect. To facilitate physiological interpretation, we apply absolute value modulation in the wiring equation 1, constraining *η <* 0 and *γ >* 0. Parameters are optimized via grid search to minimize the discrepancy between simulated and empirical network structure.

To address scalability limits, where edge computation grows exponentially with node count, we applied coarse-graining by clustering ROIs into hubs using k-means clustering (Figure S5). This preserves spatial structure while reducing variance compared to node down-sampling. Regional resolution effects are illustrated using the habenula (Figure S5). Simulations terminated at 10% edge density.

- **Whole-brain simulation:** 500 hubs (250 per hemisphere), 300 trials pre-PTZ.
- **Local-circuit simulation:** 16 regions selected from results in Figure 5C at single-cell resolution, 100 trials over 20 time steps.

#### Network formation dynamics

At each time step, edges form probabilistically based on *P*_*ij*_ (Eq. 1), balancing wiring cost 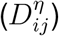 and topological reward 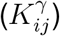. The matching index *K*_*ij*_ quantifies shared connectivity profiles, excluding self-connections and inter-node links, and is symmetric by definition^45^.

Connections are binary and static once formed. To identify optimal wiring rules, we performed 90,000 simulations across *η* = [−1.5, 0] and *γ* = [0, 1], with fine increments (0.1 and 0.05). Inputs included cell coordinates or coarse-grained node positions, with proximity serving as a heuristic reward. Simulations progressed from an unconnected graph to 25,000 edges (out of 250,000 possible), executed on the University of Melbourne’s Spartan HPC system (4 CPUs, 80 GB RAM^81^). Each simulation was repeated 300 times with randomized initial conditions to ensure robustness.

#### Evaluation of generative network models

To evaluate the biological plausibility of simulated networks, we calibrated the generative model parameters *η* and *γ* using a probabilistic wiring framework and validated outputs against empirical data. Rather than aiming for exact replication, which is impractical due to individual variability^86^, we assessed network fidelity via an “energy” score, defined as the maximum Kolmogorov–Smirnov (KS) divergence across four graph metrics: node degree (*KS*_*d*_), clustering coefficient (*KS*_*c*_), betweenness centrality (*KS*_*b*_), and edge length (*KS*_*e*_) ^45,46,63,87^.

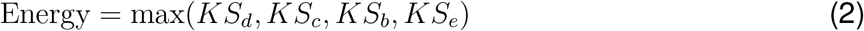

Model fit was defined by the least favorable mismatch across graph metrics, offering a conservative and robust assessment of topological similarity. Lower energy values reflected closer alignment between simulated and empirical networks, preserving key features such as local efficiency, hub resilience, and spatial embedding^88^. To account for stochastic variability, 300 parallel simulations were performed per fish and PTZ stage on the University of Melbourne HPC^81^. A 2D regression surface was then fitted to the resulting energy landscape, with the global minimum of the interpolated distribution designated as the best-fit parameter pair for each condition (Figure S5).

#### Network fingerprint characterization

Each zebrafish’s energy landscape was treated as a network “fingerprint,” capturing the divergence between simulated and empirical connectivity across a two-dimensional parameter space defined by *η* and *γ*. These landscapes encode the aggregate Kolmogorov–Smirnov (KS) distance across graph theory metrics, reflecting topological deviation from empirical data. Similarity between landscapes implies shared structural features, even if the underlying networks differ in detail, enabling comparative analysis of network organization across genotypes, regions, and PTZ stages.

#### Dimensionality reduction and classification

To extract dominant patterns, principal component analysis (PCA) was applied to a 70 × 320 matrix comprising 35 optimal *η*–*γ* pairs across 20 temporal stages and 16 anatomical regions. Each row represented a specimen, and each column a fingerprint-derived feature, with an accompanying classification vector mapping features to anatomical regions (Figure S6).

Partial Least Squares Discriminant Analysis (PLS-DA) was used to classify genotypes (Figure S7), seizure severity (Figure S7), and PTZ stages (Figure S7). Input features consisted of full simulation-derived fingerprints from 16 regions, each constructed from 100 × 100 simulations spanning the *η*–*γ* space (10,000 points per fingerprint). This enabled longitudinal tracking of network dynamics across developmental stages and genotypes.

For genotype classification at the pre-PTZ stage, a 160, 000 × 35 matrix was generated (16 regions × 10,000 points × 35 specimens), with labels distinguishing WT and *scn1lab*^−/−^ fish. Seizure severity was binned into high and low categories, with additional bin sizes (2–5) explored (Figure S7). Although finer binning reduced predictive accuracy due to limited sample sizes, regional contributions to classification remained robust.

To classify PTZ-induced developmental stages, mean network correlation was used as a proxy for excitability. Correlation values were segmented into five bins, each mapped to corresponding *η*–*γ* coordinates. Values below 0.1 indicated baseline activity, while values above 0.49 reflected frequent seizures and heightened synchronization, consistent with ictogenesis progression (Figure S7).

### Quantification and statistical analysis

All statistical analyses were performed in MATLAB R2022b and GraphPad Prism 10.0. No formal sample size estimation was performed; sample sizes were based on prior studies and standard practices in the field. Animals were randomly assigned to experimental groups, and no data or subjects were excluded unless specified in the figure legends. No stratification methods were applied.

#### Student’s t-test

In Figures 1C and 1D, Student’s t-tests were performed to compare genotypes based on total seizure counts per animal and inter-seizure intervals. Seventeen WT and eighteen *scn1lab*^−/−^ mutant siblings were analyzed. Data are presented as group means.

In Figure 1E, seizure counts were compared across genotypes at 13 distinct time points using Student’s t-tests with Bonferroni correction (N = 13). Statistical significance was defined as a corrected p *<* 0.05.

#### One-way analysis of variance with Šidák’s correction

One-way ANOVA with Šidák’s post-hoc test was used for regional comparisons in Figure S1. This approach controls the family-wise error rate while maintaining statistical power across multiple regional contrasts. This analysis enabled robust identification of region-specific phenotypic differences between WT and *scn1lab*^−/−^ genotypes.

#### Mann-Whitney U test

To evaluate genotype-specific differences without a multicomparison required, we applied the Mann—Whitney U test, a nonparametric alternative to the independent samples *t*-test. We compared the metric distributions between genotypes using MATLAB’s `ranksum` function, which tests for differences in central tendency without assuming normality.

For voxel-wise comparison of ROI density between WT and *scn1lab*^−/−^ brains, we applied the Mann—Whitney U test to each 6 × 6 × 6 *µm*^3^ voxel across the brain volume in Figure 3B. This non-parametric test was chosen to accommodate potential differences in distribution shape and variance between genotypes. Voxels with p *<* 0.05 were considered significantly different.

#### Repeated measures analysis of variance (rANOVA)

To evaluate genotype-dependent differences across multiple longitudinal time points, we performed a repeated measures analysis of variance using MATLAB’s `fitrm` and `ranova` functions. The data were structured in wide format, with each row representing a subject and each column corresponding to a distinct time point. Genotype was treated as a between-subject factor, while time was modeled as a within-subject factor. The repeated measures model was specified using `fitrm`, followed by within-subject ANOVA via `ranova`, enabling formal testing of genotype × time interactions. To identify specific time points at which genotypes diverged, we conducted post hoc pairwise comparisons using Bonferroni correction for multiple testing, implemented via the `multcompare` function on the model’s `stats` output. This approach allowed us to localize the onset and persistence of genotype effects across the temporal profile while controlling for family-wise error rate.

We applied repeated measures ANOVA to assess genotype-dependent effects of PTZ over time across graph (Figure 5; Figure S4) and broader network properties (Figure 4; Figures S3, S4 including functional connectivity (Figure 3F) and the skewness of its correlation histogram (Figure S1C). Genotype-specific differences in GNN model parameters were similarly evaluated using repeated measures ANOVA (Figure 4, 6; Figures S5, S7).

#### Effect size estimation

To quantify genotype-related effects (Figure 5B), Cohen’s *f* was derived from repeated-measures ANOVA via the transformation 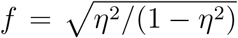, where *η*^2^ denotes the proportion of variance explained. Cohen’s *d* was used to quantify standardized differences in continuous parameters between genotypes (Figure 6E and Figure 7A). It was calculated as 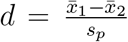, where 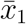 and 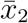 are group means for WT and *scn1lab*^−/−^ fish, respectively, and *s*_*p*_ is the pooled standard deviation. This metric provides a scale-invariant estimate of effect magnitude for direct pairwise comparisons.

#### Linear mixed-effect model (LME) to identify time effect

To assess longitudinal trends across subjects in graph metrics study in Figure 5B, we implemented a Linear mixed-effects model using MATLAB’s `fitlme` function. The dataset, originally structured as a 35 × 24 matrix (subjects × time points), was reshaped into long format, with each row representing a single observation. The model specified time as a fixed effect and included both random intercepts and slopes for each subject (`Y ∼ Time + (Time|Subject)`), allowing us to capture individual variability while testing for a consistent group-level trajectory. Statistical significance of the time effect was evaluated via ANOVA and Wald tests, with *p*-values indicating whether the observed changes over time were robust across the cohort.

#### Bonferroni correction

Following repeated measures ANOVA, Bonferroni correction was applied as a post-hoc method to adjust for multiple pairwise comparisons across conditions. This approach controls the family-wise error rate by dividing the significance threshold (*α*) by the number of comparisons, thereby reducing the likelihood of Type I errors. Pairwise contrasts were evaluated only when the overall ANOVA yielded a significant effect. Corrected *p*-values were reported, and significance was defined at the adjusted threshold.

## Additional resources

Figure S1 was created using BioRender.com to illustrate the experimental workflow and anatomical reference framework.

## RESOURCE AVAILABILITY

### Lead contact

Requests for further information and resources should be directed to and will be fulfilled by the lead contact, Wei Qin.

### Materials availability

No novel genetic constructs, mutants, or transgenic lines were generated.

### Data and code availability

- All original code has been deposited at Github (https://github.com/qinwayne/scn1lab) and is publicly available as of the date of publication.
- Any additional information required to reanalyze the data reported in this paper is available from the lead contact upon request.

## ACKNOWLEDGMENTS

Support was provided by a Simons Foundation Research Award (625793), two ARC Discovery Project Grants (DP220103812 and DP230102614), and an NHMRC Investigator Grant (2027072) to E.K.S. The research reported in this publication was supported by the National Institute of Neurological Disorders and Stroke of the National Institutes of Health under Award Number R01NS118406 to E.K.S. The content is solely the responsibility of the authors and does not necessarily represent the official views of the National Institutes of Health. Support was also provided by an ARC DECRA award (DE230100972) and an NHMRC Ideas Grant (2012140) to I.A.F. M.W. was supported by a University of Queensland RTP scholarship.

## AUTHOR CONTRIBUTIONS

Conceptualization, E.K.S. and W.Q.; methodology, W.Q., J.B., and E.K.S.; investigation, W.Q. and J.B.; writing: original draft, W.Q. and E.K.S.; writing: review & editing, W.Q., E.K.S., M.W., S.J.S, M.M., and I.A.F.; funding acquisition, E.K.S. and I.A.F.; resources, E.K.S.; supervision, E.K.S.

## DECLARATION OF INTERESTS

The authors declare no competing interests.

## SUPPLEMENTAL INFORMATION INDEX

Figures S1-S7 and their legends in a PDF

Video 1. Behavioral manifestations observed in zebrafish during seizure development.

Video 2. An animated video showing comparison between SyN and ROI density over different planes.

Video 3. An animated video illustrates the temporal dynamics of correlation strength across ipsilateral and contralateral connections, comparing region-to-region and region-to-hemisphere interactions. Similar to S3

## Notes

### Competing Interest Statement

The authors have declared no competing interest.

## References

1. Ritter-Makinson, S., Clemente-Perez, A., Higashikubo, B., Cho, F.S., Holden, S.S., Bennett, E., Chkhaidze, A., Rooda, O.H.E., Cornet, M.C., Hoebeek, F.E. et al. (2019). Augmented reticular thalamic bursting and seizures in scn1a-dravet syndrome. Cell reports 26, 54–64.

2. Holmes, G.L. (2016). Effect of seizures on the developing brain and cognition. In Seminars in pediatric neurology vol. 23. Elsevier pp. 120–126.

3. Laxer, K.D., Trinka, E., Hirsch, L.J., Cendes, F., Langfitt, J., Delanty, N., Resnick, T., and Benbadis, S.R. (2014). The consequences of refractory epilepsy and its treatment. Epilepsy & behavior 37, 59–70.

4. Tomson, T., Nashef, L., and Ryvlin, P. (2008). Sudden unexpected death in epilepsy: current knowledge and future directions. The Lancet Neurology 7, 1021–1031.

5. Dalic, L., and Cook, M.J. (2016). Managing drug-resistant epilepsy: challenges and solutions. Neuropsychiatric disease and treatment pp. 2605–2616.

6. Jirsa, V.K., Proix, T., Perdikis, D., Woodman, M.M., Wang, H., Gonzalez-Martinez, J., Bernard, C., Bénar, C., Guye, M., Chauvel, P. et al. (2017). The virtual epileptic patient: individualized whole-brain models of epilepsy spread. Neuroimage 145, 377–388.

7. Wendling, F., Benquet, P., Bartolomei, F., and Jirsa, V. (2016). Computational models of epileptiform activity. Journal of neuroscience methods 260, 233–251.

8. Proix, T., Bartolomei, F., Guye, M., and Jirsa, V.K. (2017). Individual brain structure and modelling predict seizure propagation. Brain 140, 641–654.

9. Lytton, W.W. (2008). Computer modelling of epilepsy. Nature Reviews Neuroscience 9, 626–637.

10. Goodfellow, M., Rummel, C., Abela, E., Richardson, M.P., Schindler, K., and Terry, J.R. (2016). Estimation of brain network ictogenicity predicts outcome from epilepsy surgery. Scientific reports 6, 29215.

11. Sinha, N., Dauwels, J., Kaiser, M., Cash, S.S., Brandon Westover, M., Wang, Y., and Taylor, P.N. (2017). Predicting neurosurgical outcomes in focal epilepsy patients using computational modelling. Brain 140, 319–332.

12. Suffczynski, P., Da Silva, F.H.L., Parra, J., Velis, D.N., Bouwman, B.M., Van Rijn, C.M., Van Hese, P., Boon, P., Khosravani, H., Derchansky, M. et al. (2006). Dynamics of epileptic phenomena determined from statistics of ictal transitions. IEEE transactions on biomedical engineering 53, 524–532.

13. Vanwalleghem, G.C., Ahrens, M.B., and Scott, E.K. (2018). Integrative whole-brain neuroscience in larval zebrafish. Current opinion in neurobiology 50, 136–145.

14. Hasani, H., Sun, J., Zhu, S.I., Rong, Q., Willomitzer, F., Amor, R., McConnell, G., Cossairt, O., and Goodhill, G.J. (2023). Whole-brain imaging of freely-moving zebrafish. Frontiers in neuroscience 17, 1127574.

15. Doszyn, O., Dulski, T., and Zmorzynska, J. (2024). Diving into the zebrafish brain: exploring neuroscience frontiers with genetic tools, imaging techniques, and behavioral insights. Frontiers in Molecular Neuroscience 17, 1358844.

16. Wullimann, M., Rupp, B., Reichert, H., Wullimann, M., Rupp, B., and Reichert, H. (1996). Functional anatomy of the zebrafish brain: a comparative evaluation. Neuroanatomy of the Zebrafish Brain: A Topological Atlas pp. 89–101.

17. Baraban, S.C., Taylor, M., Castro, P., and Baier, H. (2005). Pentylenetetrazole induced changes in zebrafish behavior, neural activity and c-fos expression. Neuroscience 131, 759–768.

18. D’Iglio, C., Famulari, S., Capparucci, F., Gervasi, C., Cuzzocrea, S., Spanò, N., and Di Paola, D. (2023). Toxic effects of gemcitabine and paclitaxel combination: Chemotherapy drugs exposure in zebrafish. Toxics 11, 544.

19. Niemeyer, J.E., Gadamsetty, P., Chun, C., Sylvester, S., Lucas, J.P., Ma, H., Schwartz, T.H., and Aksay, E.R. (2022). Seizures initiate in zones of relative hyperexcitation in a zebrafish epilepsy model. Brain 145, 2347–2360.

20. Yaksi, E., Jamali, A., Diaz Verdugo, C., and Jurisch-Yaksi, N. (2021). Past, present and future of zebrafish in epilepsy research. The FEBS journal 288, 7243–7255.

21. Aksay, E., Olasagasti, I., Mensh, B.D., Baker, R., Goldman, M.S., and Tank, D.W. (2007). Functional dissection of circuitry in a neural integrator. Nature neuroscience 10, 494–504.

22. Griffin, A., Hamling, K.R., Knupp, K., Hong, S., Lee, L.P., and Baraban, S.C. (2017). Clemizole and modulators of serotonin signalling suppress seizures in dravet syndrome. Brain 140, 669–683.

23. Baxendale, S., Holdsworth, C.J., Meza Santoscoy, P.L., Harrison, M.R., Fox, J., Parkin, C.A., Ingham, P.W., and Cunliffe, V.T. (2012). Identification of compounds with anti-convulsant properties in a zebrafish model of epileptic seizures. Disease models & mechanisms 5, 773–784.

24. Ahrens, M.B., Orger, M.B., Robson, D.N., Li, J.M., and Keller, P.J. (2013). Whole-brain functional imaging at cellular resolution using light-sheet microscopy. Nature methods 10, 413–420.

25. Favre-Bulle, I.A., Vanwalleghem, G., Taylor, M.A., Rubinsztein-Dunlop, H., and Scott, E.K. (2018). Cellular-resolution imaging of vestibular processing across the larval zebrafish brain. Current Biology 28, 3711–3722.

26. Afrikanova, T., Serruys, A.S.K., Buenafe, O.E., Clinckers, R., Smolders, I., de Witte, P.A., Crawford, A.D., and Esguerra, C.V. (2013). Validation of the zebrafish pentylenetetrazol seizure model: locomotor versus electrographic responses to antiepileptic drugs. PloS one 8, e54166.

27. Constantin, L., Poulsen, R.E., Scholz, L.A., Favre-Bulle, I.A., Taylor, M.A., Sun, B., Goodhill, G.J., Vanwalleghem, G.C., and Scott, E.K. (2020). Altered brain-wide auditory networks in a zebrafish model of fragile x syndrome. BMC biology 18, 1–17.

28. Mancienne, T., Marquez-Legorreta, E., Wilde, M., Piber, M., Favre-Bulle, I., Vanwalleghem, G., and Scott, E.K. (2021). Contributions of luminance and motion to visual escape and habituation in larval zebrafish. Frontiers in neural circuits 15, 748535.

29. Pachitariu, M., Stringer, C., Schröder, S., Dipoppa, M., Rossi, L.F., Carandini, M., and Harris, K.D. (2016). Suite2p: beyond 10,000 neurons with standard two-photon microscopy. BioRxiv pp. 061507.

30. Randlett, O., Wee, C.L., Naumann, E.A., Nnaemeka, O., Schoppik, D., Fitzgerald, J.E., Portugues, R., Lacoste, A.M., Riegler, C., Engert, F. et al. (2015). Whole-brain activity mapping onto a zebrafish brain atlas. Nature methods 12, 1039–1046.

31. Tustison, N.J., Cook, P.A., Holbrook, A.J., Johnson, H.J., Muschelli, J., Devenyi, G.A., Duda, J.T., Das, S.R., Cullen, N.C., Gillen, D.L. et al. (2021). The antsx ecosystem for quantitative biological and medical imaging. Scientific reports 11, 9068.

32. Marquez-Legorreta, E., Constantin, L., Piber, M., Favre-Bulle, I.A., Taylor, M.A., Blevins, A.S., Giacomotto, J., Bassett, D.S., Vanwalleghem, G.C., and Scott, E.K. (2022). Brain-wide visual habituation networks in wild type and fmr1 zebrafish. Nature Communications 13, 895.

33. Wilde, M., Ghanbari, A., Mancienne, T., Moran, A., Poulsen, R.E., Constantin, L., Lee, C., Scholz, L.A., Arnold, J., Qin, W. et al. (2024). Brain-wide circuitry underlying altered auditory habituation in zebrafish models of autism. bioRxiv.

34. Hong, F.L. (2016). Optical frequency standards for time and length applications. Measurement Science and Technology 28, 012002.

35. Avants, B.B., Tustison, N.J., Stauffer, M., Song, G., Wu, B., and Gee, J.C. (2014). The insight toolkit image registration framework. Frontiers in neuroinformatics 8, 44.

36. Avants, B.B., Epstein, C.L., Grossman, M., and Gee, J.C. (2008). Symmetric diffeomorphic image registration with cross-correlation: evaluating automated labeling of elderly and neurodegenerative brain. Medical image analysis 12, 26–41.

37. Diaz Verdugo, C., Myren-Svelstad, S., Aydin, E., Van Hoeymissen, E., Deneubourg, C., Vanderhaeghe, S., Vancraeynest, J., Pelgrims, R., Cosacak, M.I., Muto, A. et al. (2019). Glia-neuron interactions underlie state transitions to generalized seizures. Nature communications 10, 3830.

38. Palva, J.M., and Palva, S. (2014). The correlation of the neuronal long-range temporal correlations, avalanche dynamics with the behavioral scaling laws and interindividual variability. Criticality in Neural Systems pp. 105–126.

39. Shen, B., Wei, H., Wen, Y., Geng, Y., Yang, T., Chen, Z., Dong, S., Gao, Y., Li, T., Sun, L. et al. (2025). Optimized in vivo two-photon imaging reveals the essential role of the contralateral eye in functional optic nerve regeneration in zebrafish larvae. Eye and Vision 12, 34.

40. Rubinov, M., Kötter, R., Hagmann, P., and Sporns, O. (2009). Brain connectivity toolbox: a collection of complex network measurements and brain connectivity datasets. NeuroImage 47, S169.

41. Wang, J., Zuo, X., and He, Y. (2010). Graph-based network analysis of resting-state functional mri. Frontiers in systems neuroscience 4, 1419.

42. Liao, W., Zhang, Z., Pan, Z., Mantini, D., Ding, J., Duan, X., Luo, C., Lu, G., and Chen, H. (2010). Altered functional connectivity and small-world in mesial temporal lobe epilepsy. PloS one 5, e8525.

43. Bassett, D.S., and Bullmore, E.T. (2009). Human brain networks in health and disease. Current opinion in neurology 22, 340–347.

44. Bialonski, S., and Lehnertz, K. (2013). Assortative mixing in functional brain networks during epileptic seizures. Chaos: An Interdisciplinary Journal of Nonlinear Science 23.

45. Akarca, D., Vértes, P.E., Bullmore, E.T., and Astle, D.E. (2021). A generative network model of neurodevelopmental diversity in structural brain organization. Nature communications 12, 4216.

46. Betzel, R.F., Avena-Koenigsberger, A., Goñi, J., He, Y., De Reus, M.A., Griffa, A., Vértes, P.E., Mišic, B., Thiran, J.P., Hagmann, P. et al. (2016). Generative models of the human connectome. Neuroimage 124, 1054–1064.

47. Vértes, P.E., Alexander-Bloch, A.F., Gogtay, N., Giedd, J.N., Rapoport, J.L., and Bullmore, E.T. (2012). Simple models of human brain functional networks. Proceedings of the National Academy of Sciences 109, 5868–5873.

48. Liu, Y., Seguin, C., Mansour, S., Oldham, S., Betzel, R., Di Biase, M.A., and Zalesky, A. (2023). Parameter estimation for connectome generative models: Accuracy, reliability, and a fast parameter fitting method. Neuroimage 270, 119962.

49. Svob Strac, D., Pivac, N., Smolders, I.J., Fogel, W.A., De Deurwaerdere, P., and Di Giovanni, G. (2016). Monoaminergic mechanisms in epilepsy may offer innovative therapeutic opportunity for monoaminergic multi-target drugs. Frontiers in neuroscience 10, 492.

50. Fishell, G., and Rudy, B. (2011). Mechanisms of inhibition within the telencephalon:”where the wild things are”. Annual review of neuroscience 34, 535–567.

51. Uysal, S. (2023). Functional neuroanatomy and clinical neuroscience: foundations for understanding disorders of cognition and behavior. Oxford University Press.

52. Marafiga, J.R., Pasquetti, M.V., and Calcagnotto, M.E. (2021). Gabaergic interneurons in epilepsy: More than a simple change in inhibition. Epilepsy & Behavior 121, 106935.

53. Deco, G., Kringelbach, M.L., Jirsa, V.K., and Ritter, P. (2017). The dynamics of resting fluctuations in the brain: metastability and its dynamical cortical core. Scientific reports 7, 3095.

54. Lin, X.Y., Cui, Y., Wang, L., and Chen, W. (2019). Chronic exercise buffers the cognitive dysfunction and decreases the susceptibility to seizures in ptz-treated rats. Epilepsy & Behavior 98, 173–187.

55. Egger, M., Luo, W., Cruz-Ochoa, N., Lukacsovich, D., Varga, C., Que, L., Maloveczky, G., Winterer, J., Kaur, R., Lukacsovich, T. et al. (2023). Commissural dentate granule cell projections and their rapid formation in the adult brain. PNAS nexus 2, pgad088.

56. Schindler, K., Leung, H., Lehnertz, K., and Elger, C.E. (2007). How generalised are secondarily “generalised” tonic–clonic seizures? Journal of Neurology, Neurosurgery & Psychiatry 78, 993–996.

57. Gerster, M., Berner, R., Sawicki, J., Zakharova, A., Škoch, A., Hlinka, J., Lehnertz, K., and Schöll, E. (2020). Fitzhugh–nagumo oscillators on complex networks mimic epileptic-seizure-related synchronization phenomena. Chaos: An Interdisciplinary Journal of Nonlinear Science 30.

58. Zhou, D., Stanley, H.E., D’Agostino, G., and Scala, A. (2012). Assortativity decreases the robustness of interdependent networks. Physical Review E—Statistical, Nonlinear, and Soft Matter Physics 86, 066103.

59. D’Agostino, G., Scala, A., Zlatic, V., and Caldarelli, G. (2011). Assortativity effects on diffusion-like processes in scale-free networks. arXiv preprint arXiv:1105.3574.

60. Pedersen, M., Omidvarnia, A.H., Walz, J.M., and Jackson, G.D. (2015). Increased segregation of brain networks in focal epilepsy: An fmri graph theory finding. NeuroImage: Clinical 8, 536–542.

61. Sporns, O. (2013). Network attributes for segregation and integration in the human brain. Current opinion in neurobiology 23, 162–171.

62. Deco, G., Jirsa, V.K., and McIntosh, A.R. (2011). Emerging concepts for the dynamical organization of resting-state activity in the brain. Nature reviews neuroscience 12, 43–56.

63. Bassett, D.S., and Bullmore, E.T. (2017). Small-world brain networks revisited. The Neuroscientist 23, 499–516.

64. Bassett, D.S., Bullmore, E., Verchinski, B.A., Mattay, V.S., Weinberger, D.R., and Meyer-Lindenberg, A. (2008). Hierarchical organization of human cortical networks in health and schizophrenia. Journal of Neuroscience 28, 9239–9248.

65. Stam, C.J. (2014). Modern network science of neurological disorders. Nature Reviews Neuroscience 15, 683–695.

66. Fornito, A., Zalesky, A., and Breakspear, M. (2015). The connectomics of brain disorders. Nature Reviews Neuroscience 16, 159–172.

67. Filippi, A., Mueller, T., and Driever, W. (2014). vglut2 and gad expression reveal distinct patterns of dual gabaergic versus glutamatergic cotransmitter phenotypes of dopaminergic and noradrenergic neurons in the zebrafish brain. Journal of Comparative Neurology 522, 2019–2037.

68. Macdonald, R.L., and Barker, J.L. (1978). Specific antagonism of gaba-mediated postsynaptic inhibition in cultured mammalian spinal cord neurons: a common mode of convulsant action. Neurology 28, 325–325.

69. Valassina, N., Brusco, S., Salamone, A., Serra, L., Luoni, M., Giannelli, S., Bido, S., Massimino, L., Ungaro, F., Mazzara, P.G. et al. (2022). Scn1a gene reactivation after symptom onset rescues pathological phenotypes in a mouse model of dravet syndrome. Nature communications 13, 161.

70. Depienne, C., Trouillard, O., Saint-Martin, C., Gourfinkel-An, I., Bouteiller, D., Carpentier, W., Keren, B., Abert, B., Gautier, A., Baulac, S. et al. (2009). Spectrum of scn1a gene mutations associated with dravet syndrome: analysis of 333 patients. Journal of medical genetics 46, 183–191.

71. Griffin, A., Hamling, K.R., Hong, S., Anvar, M., Lee, L.P., and Baraban, S.C. (2018). Pre-clinical animal models for dravet syndrome: seizure phenotypes, comorbidities and drug screening. Frontiers in pharmacology 9, 573.

72. Brenet, A., Hassan-Abdi, R., Somkhit, J., Yanicostas, C., and Soussi-Yanicostas, N. (2019). Defective excitatory/inhibitory synaptic balance and increased neuron apoptosis in a zebrafish model of dravet syndrome. Cells 8, 1199.

73. Liu, J., and Baraban, S.C. (2019). Network properties revealed during multi-scale calcium imaging of seizure activity in zebrafish. eneuro 6.

74. Zhang, Y., Heylen, L., Partoens, M., Mills, J.D., Kaminski, R.M., Godard, P., Gillard, M., de Witte, P.A., and Siekierska, A. (2022). Connectivity mapping using a novel sv2a loss-of-function zebrafish epilepsy model as a powerful strategy for anti-epileptic drug discovery. Frontiers in Molecular Neuroscience 15, 881933.

75. Wong, J.C., Dutton, S.B., Collins, S.D., Schachter, S., and Escayg, A. (2016). Huperzine a provides robust and sustained protection against induced seizures in scn1a mutant mice. Frontiers in Pharmacology 7, 357.

76. Das, A., Zhu, B., Xie, Y., Zeng, L., Pham, A.T., Neumann, J.C., Safrina, O., Benavides, D.R., MacGregor, G.R., Schutte, S.S. et al. (2021). Interneuron dysfunction in a new mouse model of scn1a gefs+. ENeuro 8.

77. Parihar, R., and Ganesh, S. (2013). The scn1a gene variants and epileptic encephalopathies. Journal of human genetics 58, 573–580.

78. Fan, D., Wu, H., Luan, G., and Wang, Q. (2023). The distribution and heterogeneity of excitability in focal epileptic network potentially contribute to the seizure propagation. Frontiers in Psychiatry 14, 1137704.

79. Marques, J.C., Li, M., Schaak, D., Robson, D.N., and Li, J.M. (2020). Internal state dynamics shape brainwide activity and foraging behaviour. Nature 577, 239–243.

80. Taylor, M.A., Vanwalleghem, G.C., Favre-Bulle, I.A., and Scott, E.K. (2018). Diffuse lightsheet microscopy for stripe-free calcium imaging of neural populations. Journal of biophotonics 11, e201800088.

81. Research Computing Services, U.o.M. (2025). Spartan high-performance computing system. URL: https://dashboard.hpc.unimelb.edu.au/ accessed: April 28, 2025.

82. Vanwalleghem, G., Constantin, L., and Scott, E.K. (2021). Calcium imaging and the curse of negativity. Frontiers in neural circuits 14, 607391.

83. Stafstrom, C.E., and Carmant, L. (2015). Seizures and epilepsy: an overview for neuroscientists. Cold Spring Harbor perspectives in medicine 5, a022426.

84. Maslov, S., and Sneppen, K. (2002). Specificity and stability in topology of protein networks. Science 296, 910–913.

85. Váša, F., and Mišić, B. (2022). Null models in network neuroscience. Nature Reviews Neuroscience 23, 493–504.

86. Honey, C.J., Sporns, O., Cammoun, L., Gigandet, X., Thiran, J.P., Meuli, R., and Hagmann, P. (2009). Predicting human resting-state functional connectivity from structural connectivity. Proceedings of the National Academy of Sciences 106, 2035–2040.

87. Betzel, R.F., and Bassett, D.S. (2017). Multi-scale brain networks. Neuroimage 160, 73–83.

88. Rubinov, M., and Sporns, O. (2010). Complex network measures of brain connectivity: uses and interpretations. Neuroimage 52, 1059–1069.

