## Supplementary Information for "Whole-brain cellular-resolution functional network properties of seizure susceptibility"

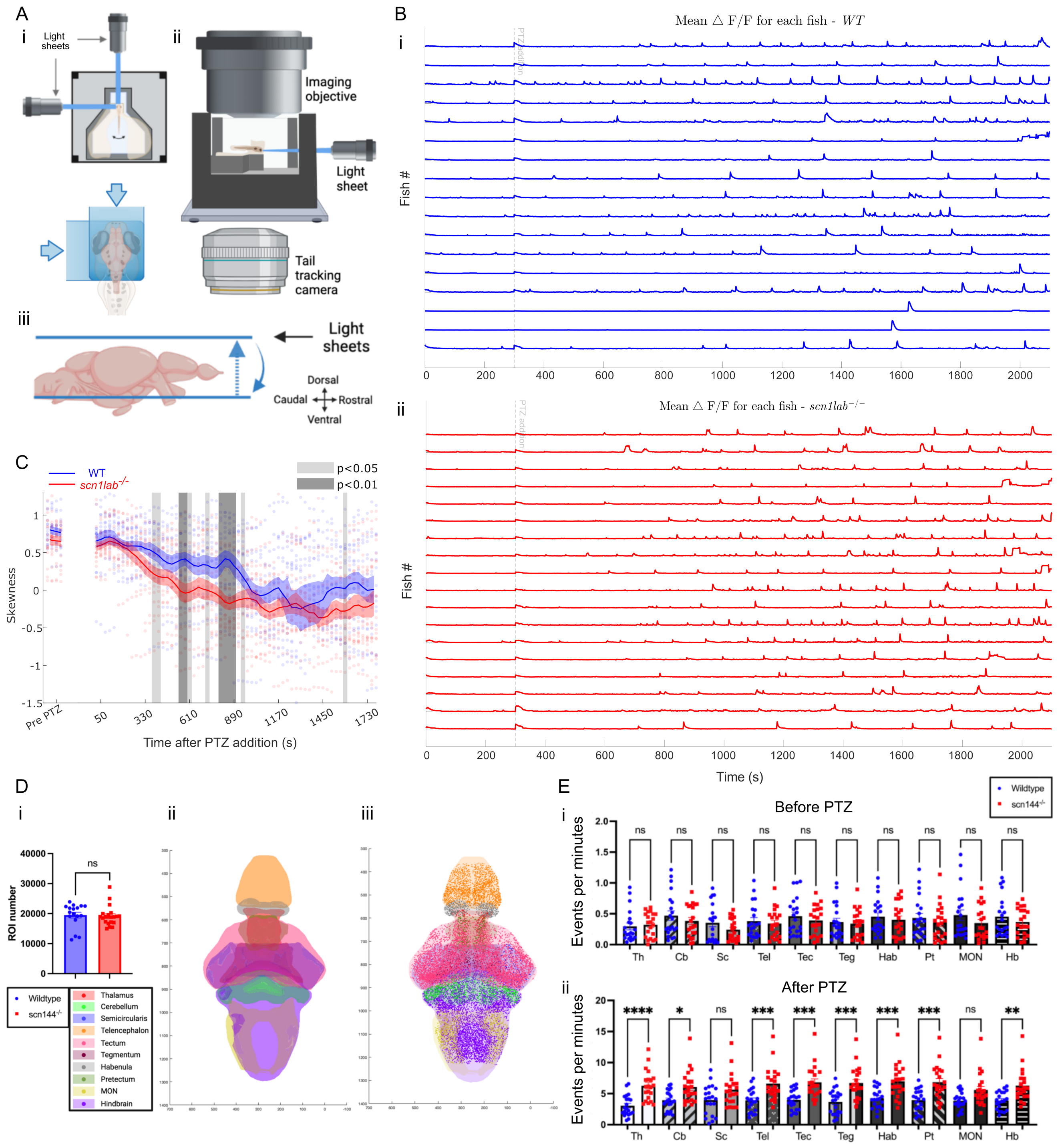

A
i
Light sheets
ii
Imaging objective
Light sheet
Tail tracking camera
iii
Light sheets
Dorsal
Caudal
Rostral
Ventral
B
i
Mean △ F/F for each fish - WT
Fish #
ii
Mean △ F/F for each fish - scn1lab-/-
Time (s)
C
WT
scn1lab-/-
p<0.05
p<0.01
Skewness
Pre PTZ
Time after PTZ addition (s)
D
i
ii
iii
ROI number
ns
Wildtype
scn144-/-
Thalamus
Cerebellum
Semicircularis
Telencephalon
Tectum
Tegmentum
Habenula
Pretectum
MON
Hindbrain
E
i
Before PTZ
Events per minutes
Wildtype
scn144-/-
Th
Cb
Sc
Tel
Tec
Teg
Hab
Pt
MON
Hb
ii
After PTZ


Figure S1

#### Figure S1. Calcium imaging and segmentation

(A) Schematic of calcium experiments. (i) Top view of imaging chamber showing the two light sheets used for imaging, one from the front of the fish and one from the side. (ii) Side view of imaging chamber showing the orthogonal imaging objective focused on the brain, and another camera below used for tail tracking. (iii) Representations of the positioning of the light sheet relative to the zebrafish brain. Each sheet overlaps in X and Y dimensions (top) and scans together through Z in 10 $\mu m$ increments to form 25 slices encapsulating a volumetric 3D scan of the brain. The bottom plane is positioned underneath the tectum, with the top plane ending immediately above the brain.
(B) Mean $\Delta F/F$. (i) Offset whole-brain $\Delta F/F$ traces for all WT larvae. Each line represents an individual larva. (ii) Offset whole-brain $\Delta F/F$ traces for all *scn1lab*$^{-/-}$ larvae. Each line represents an individual larva.
(C) Skewness of the correlation distributions like those in Figure 3E, shown for narrow time windows across the experiment, with significant differences highlighted. Blue indicates the WT group, while red represents the *scn1lab*$^{-/-}$ group. Lines indicate mean, while the shaded colored areas show Standard Error of the Mean (SEM), and shaded gray areas indicate timepoints with significance phenotypes as determined by rANOVA.
(D) ROI Selection. (i) There is no significant difference in the number of ROIs per fish between WT (blue, $n = 17$, $19480 \pm 957$) and *scn1lab*$^{-/-}$ (red, $n = 20$, $19166 \pm 739$). (ii) A dorsal view of a 3D model of zebrafish brain regions. The MON is located beneath the hindbrain, while the pretectum and semicircularis lie below the tectum. (iii) ROIs from a single fish are color-coded by brain region, where each colored spot represents a single ROI/neuron. Data are expressed as mean ± SEM. An unpaired Student's t-test shows no significant difference ($p = 0.7940$).
Brain regions analyzed include the thalamus (Th), cerebellum (Cb), semicircularis (Sc), telencephalon (Tel), tectum (Tec), tegmentum (Teg), habenula (Hab), pretectum (Pt), medial octavolateralis nucleus (MON), and hindbrain (Hb). Data are presented as mean ± SEM. Statistical analysis was conducted using one-way ANOVA with Šidák's post-hoc test for multiple comparisons (*$p < 0.0332$, **$p < 0.0021$, ***$p < 0.0002$, ****$p < 0.0001$).
(E) Regional analysis highlights differences in PTZ sensitivity across brain regions. (i) No significant differences in the baseline activity (events per minute) of individual ROIs were observed between genotypes across all brain regions. (ii) After PTZ administration, *scn1lab*$^{-/-}$ larvae exhibited significantly higher event rates compared to WT in all regions except the semicircularis (Sc) and medial octavolateralis nucleus (MON). Note the 10-fold increase in the Y-axis scale in panel B. All regions showed a significant rise in event rate from baseline to PTZ conditions (analysis not shown).

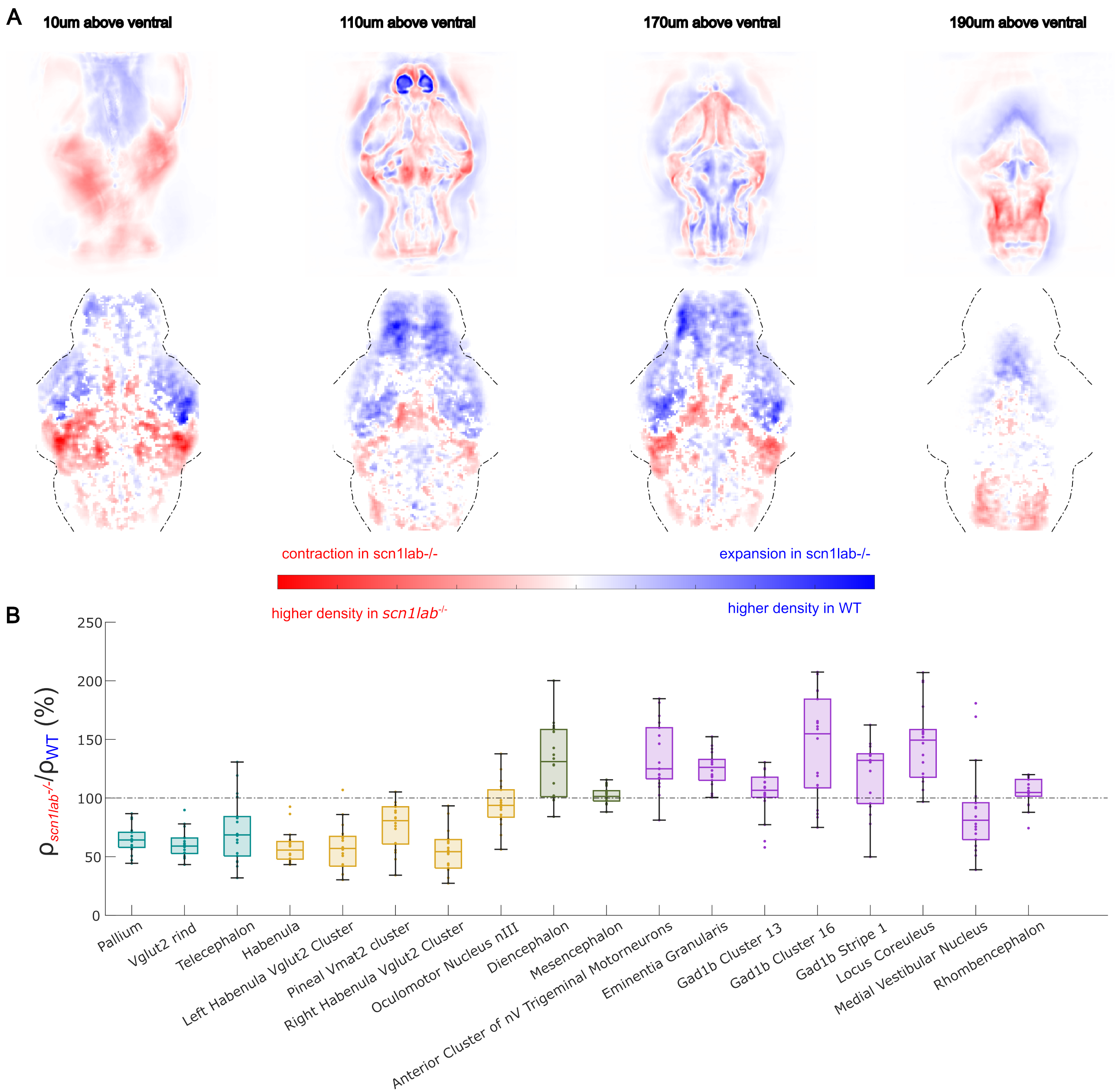
A
10um above ventral
110um above ventral
170um above ventral
190um above ventral
contraction in scn1lab-/-
expansion in scn1lab-/-
higher density in scn1lab-/-
higher density in WT
B
ρscn1lab-/-/ρWT (%)
250
200
150
100
50
0
Pallium
Vglut2 rind
Telecephalon
Habenula
Left Habenula Vglut2 Cluster
Pineal Vmat2 cluster
Right Habenula Vglut2 Cluster
Oculomotor Nucleus nIII
Diencephalon
Mesencephalon
Anterior Cluster of nV Trigeminal Motorneurons
Eminentia Granularis
Gad1b Cluster 13
Gad1b Cluster 16
Gad1b Stripe 1
Locus Coreuleus
Medial Vestibular Nucleus
Rhombencephalon

Figure S2

#### Figure S2. Brain-wide and regional changes in *scn1lab*$^{-/-}$-mutant brain morphology

(A) The SyN method confirms widespread morphological differences between mean *scn1lab*$^{-/-}$ stacks and mean WT stacks. (Top) Blue regions indicate areas where the brain is expanded in *scn1lab*$^{-/-}$, while red regions show areas of contraction via SyN analysis. (Bottom) illustrates the ROI density differences between WT and *scn1lab*$^{-/-}$. Blue represents areas with a higher density within a $6 \times 6 \times 6\ \mu m^3$ region in WT, while red indicates areas with a higher density in *scn1lab*$^{-/-}$. Voxels are colored if they have $p < 0.05$ (Mann-Whitney U test). A total of 4 planes are displayed using SyN (Top), corresponding to their ROI density planes (Bottom). The depths are given on the top of each diagram. The results obtained from the two methods are not fully aligned, likely due to distortions introduced by ANTs-based warping, as well as the limited spatial resolution inherent to the $6 \times 6 \times 6\ \mu m^3$ voxel size employed in the ROI density approach. Nonetheless, the overall trends in regional changes remain comparable. A more detailed comparison of the differences and similarities is provided in Video 2.

(B)The complete list shows the relative abundance of ROIs in smaller regions with significant differences between genotypes (Figure 3D). Statistical analysis was conducted using Student's t-test, with significance set at $p < 0.05$. Colors and arrangement correspond to the brain regions indicated in Figure 3A.

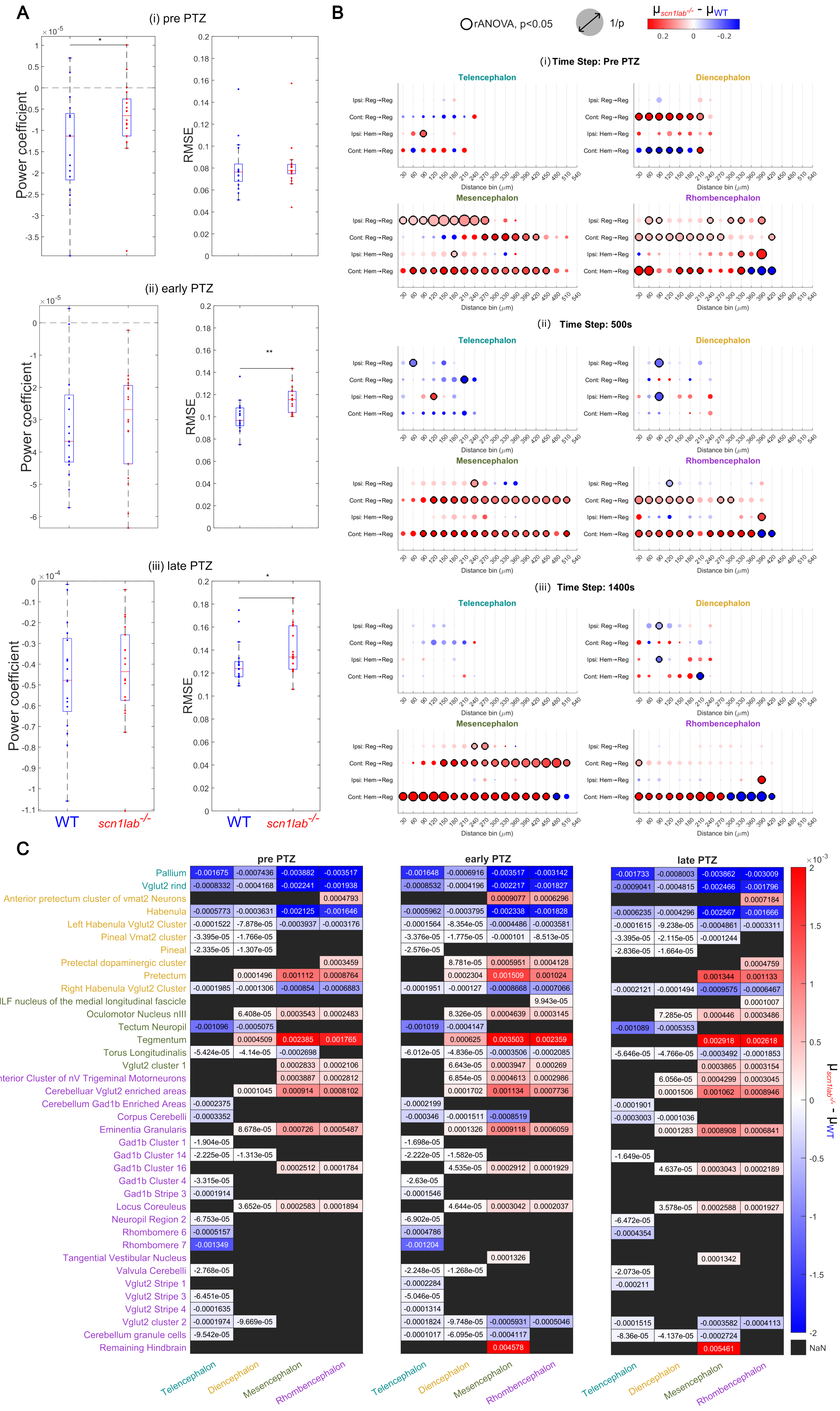

A
(i) pre PTZ
(ii) early PTZ
(iii) late PTZ
Power coefficient
RMSE
WT
scn1lab-/-
B
rANOVA, p<0.05
1/p
μscn1lab-/- - μWT
(i) Time Step: Pre PTZ
(ii) Time Step: 500s
(iii) Time Step: 1400s
Telencephalon
Diencephalon
Mesencephalon
Rhombencephalon
Ipsi: Reg→Reg
Cont: Reg→Reg
Ipsi: Hem→Reg
Cont: Hem→Reg
Distance bin (μm)
C
pre PTZ
early PTZ
late PTZ
Pallium
Vglut2 rind
Anterior pretectum cluster of vmat2 Neurons
Habenula
Left Habenula Vglut2 Cluster
Pineal Vmat2 cluster
Pineal
Pretectal dopaminergic cluster
Pretectum
Right Habenula Vglut2 Cluster
NucMLF nucleus of the medial longitudinal fascicle
Oculomotor Nucleus nIII
Tectum Neuropil
Tegmentum
Torus Longitudinalis
Vglut2 cluster 1
Anterior Cluster of nV Trigeminal Motorneurons
Cerebelluar Vglut2 enriched areas
Cerebellum Gad1b Enriched Areas
Corpus Cerebelli
Eminentia Granularis
Gad1b Cluster 1
Gad1b Cluster 14
Gad1b Cluster 16
Gad1b Cluster 4
Gad1b Stripe 3
Locus Coreuleus
Neuropil Region 2
Rhombomere 6
Rhombomere 7
Tangential Vestibular Nucleus
Valvula Cerebelli
Vglut2 Stripe 1
Vglut2 Stripe 3
Vglut2 Stripe 4
Vglut2 cluster 2
Cerebellum granule cells
Remaining Hindbrain
NaN


Figure S3

#### Figure S3. Power coefficients and correlation strengths in WT and *scn1lab*$^{-/-}$-mutant brains

(A) Linear regression of logarithmic correlation distances (0-900 $\mu m$) reveals differences between genotypes. *scn1lab*$^{-/-}$ fish show higher power coefficients, indicating stronger connectivity, especially within 200-500 $\mu m$ (Figure 4B, right). Before PTZ (i), *scn1lab*$^{-/-}$ fish exhibit significantly higher power coefficients with low RMSE, supporting a power-law distribution. No RMSE differences are observed between genotypes at this stage. RMSE increases after PTZ (ii and iii), indicating poorer fit as seizures emerge. The *scn1lab*$^{-/-}$ genotype shows a weaker fit during these stages.
(B) Correlation strength between brain regions and hemispheres was compared across ipsilateral and contralateral connections. The x-axis shows Euclidean distances between ROIs; the y-axis represents four connection types: ipsilateral region-to-region, contralateral region-to-region, ipsilateral hemisphere-to-region, and contralateral hemisphere-to-region. Data are shown at three time points: pre-PTZ, 500 s, and 1400 s. Blue indicates stronger mean correlations in WT, red in *scn1lab*$^{-/-}$. Circles denote statistically significant differences ($p < 0.05$, rANOVA). An animated movie is included in the unconventional supplementary materials (Video 3).
(C) Comparison of region-to-subregion connection counts. A sparse network was constructed using the top 10% of correlation values, capturing the strongest functional connections. Blue indicates more connections in WT, red in *scn1lab*$^{-/-}$, and black denotes no significant difference ($p > 0.05$, ANOVA). Analysis spans three PTZ stages: pre-PTZ, early-PTZ (50-900 s), and late-PTZ (900-1800 s). Color gradients reflect genotype-specific mean differences, with numerical values shown for clarity.

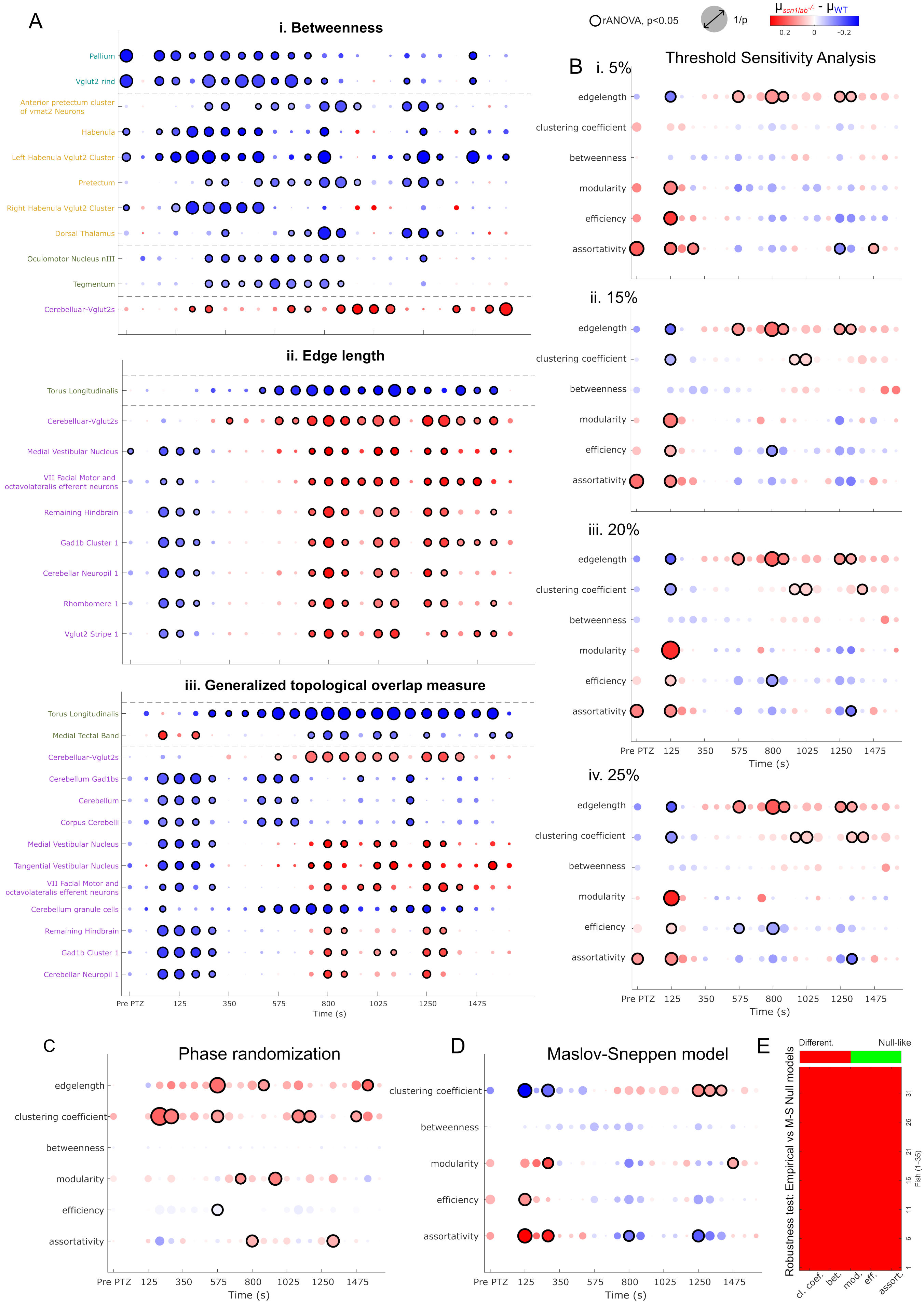

A
i. Betweenness
rANOVA, p<0.05
1/p
$\mu_{scn1lab^{-/-}} - \mu_{WT}$
0.2
0
-0.2
Pallium
Vglut2 rind
Anterior pretectum cluster of vmat2 Neurons
Habenula
Left Habenula Vglut2 Cluster
Pretectum
Right Habenula Vglut2 Cluster
Dorsal Thalamus
Oculomotor Nucleus nIII
Tegmentum
Cerebelluar-Vglut2s
ii. Edge length
Torus Longitudinalis
Cerebelluar-Vglut2s
Medial Vestibular Nucleus
VII Facial Motor and octavolateralis efferent neurons
Remaining Hindbrain
Gad1b Cluster 1
Cerebellar Neuropil 1
Rhombomere 1
Vglut2 Stripe 1
iii. Generalized topological overlap measure
Torus Longitudinalis
Medial Tectal Band
Cerebelluar-Vglut2s
Cerebellum Gad1bs
Cerebellum
Corpus Cerebelli
Medial Vestibular Nucleus
Tangential Vestibular Nucleus
VII Facial Motor and octavolateralis efferent neurons
Cerebellum granule cells
Remaining Hindbrain
Gad1b Cluster 1
Cerebellar Neuropil 1
Pre PTZ 125 350 575 800 1025 1250 1475
Time (s)
B
Threshold Sensitivity Analysis
i. 5%
ii. 15%
iii. 20%
iv. 25%
edgelength
clustering coefficient
betweenness
modularity
efficiency
assortativity
Pre PTZ 125 350 575 800 1025 1250 1475
Time (s)
C
Phase randomization
D
Maslov-Sneppen model
E
Different.
Null-like
Robustness test: Empirical vs M-S Null models
Fish (1-35)
cl. coef.
bet.
mod.
eff.
assort.


Figure S4

#### Figure S4. Node-centric metrics comparison

(A) Average values of betweenness centrality, edge length, and GTOM were computed across all cells within each subregion. Red/blue circles indicate higher means in *scn1lab*$^{-/-}$ and WT, respectively; circle size reflects confidence (1/p), and black rings denote significance ($p < 0.01$, rANOVA).
(B) Threshold Sensitivity Analysis. Genotype differences in graph metrics remain consistent across correlation thresholds (5–25%), confirming robustness of the observed effects.
(C) Phase-Randomized Null Model. Genotype-specific patterns are lost under phase randomization, indicating that the observed differences arise from biological signal rather than methodological bias.
(D) Maslov–Sneppen Null Model. 100 simulations of MS null model preserve degree distribution and yield results similar to the 10% empirical data.
(E) Null vs. Empirical Distribution Comparison. Metric distributions differ significantly between null and empirical networks, but genotype effects persist (D). This suggests that genotype differences are not independent of degree, but are embedded within broader network topology.

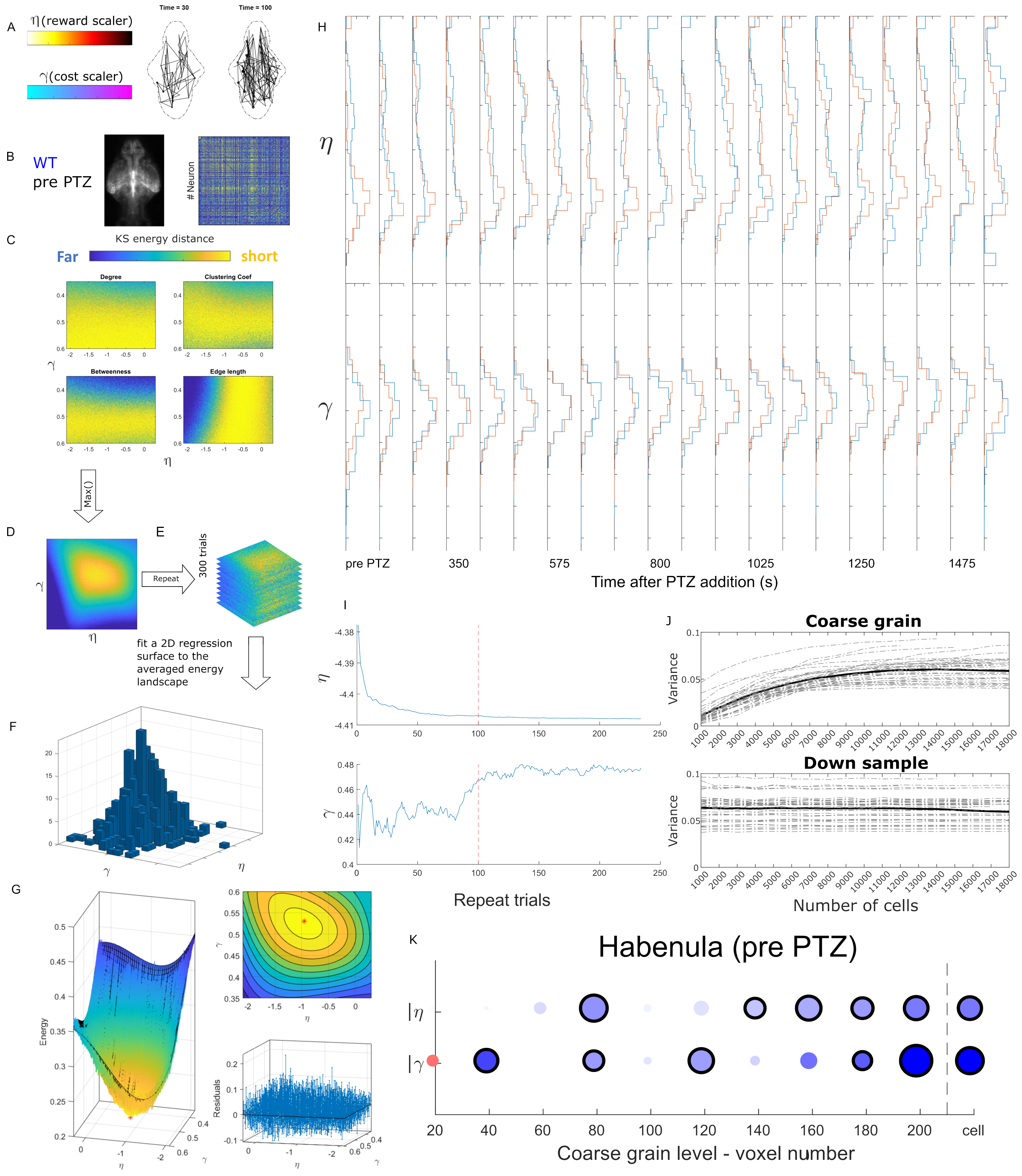

A
η(reward scaler)
γ(cost scaler)
Time = 30
Time = 100
B
WT
pre PTZ
### Neuron
C
KS energy distance
Far
short
Degree
Clustering Coef
Betweenness
Edge length
Max()
D
E
Repeat
300 trials
fit a 2D regression surface to the averaged energy landscape
F
G
Energy
Residuals
H
pre PTZ
350
575
800
1025
1250
1475
Time after PTZ addition (s)
I
Repeat trials
J
Coarse grain
Down sample
Variance
Number of cells
K
Habenula (pre PTZ)
Coarse grain level - voxel number
cell


Figure S5

#### Figure S5. GNM edge formation and evaluation workflow

(A) Edges are probabilistically formed based on paired values of $\eta$ and $\gamma$, continuing until the network matches the empirical edge count (10% of all possible connections).
(B) The top 10% of strongest correlations from the empirical dataset are used for comparison.
(C) Four graph metrics, node degree, clustering coefficient, betweenness centrality, and edge length, are computed to assess structural similarity between simulated and empirical networks.
(D) Kolmogorov–Smirnov (KS) distances quantify the maximum divergence across these metrics for each $\eta - \gamma$ pair.
(E–F) To reduce bias, simulations are repeated 300 times, and a 2D regression surface is fit to the KS results.
(G) The global minimum identifies the best-fit $\eta$ and $\gamma$ values, reflecting optimal network alignment.
(H) Histograms of $\eta$ and $\gamma$ across 20 stages show genotype-specific shifts in network configuration; blue and red lines represent WT and *scn1lab*$^{-/-}$, respectively.
(I) Convergence analysis indicates that 100 trials suffice for subregional models (red line).
(J) The variance of $\eta$ and $\gamma$ distributions through coarse graining stabilizes after 10,000 cells, whereas downsampling preserves key features with the same reduced node count.
(K) Resolution analysis in the habenula (pre-PTZ) reveals that at least 140 voxels are needed to detect genotype effects, which are clearly resolved at single-cell scale (see Figure 7B for full genotype comparisons).

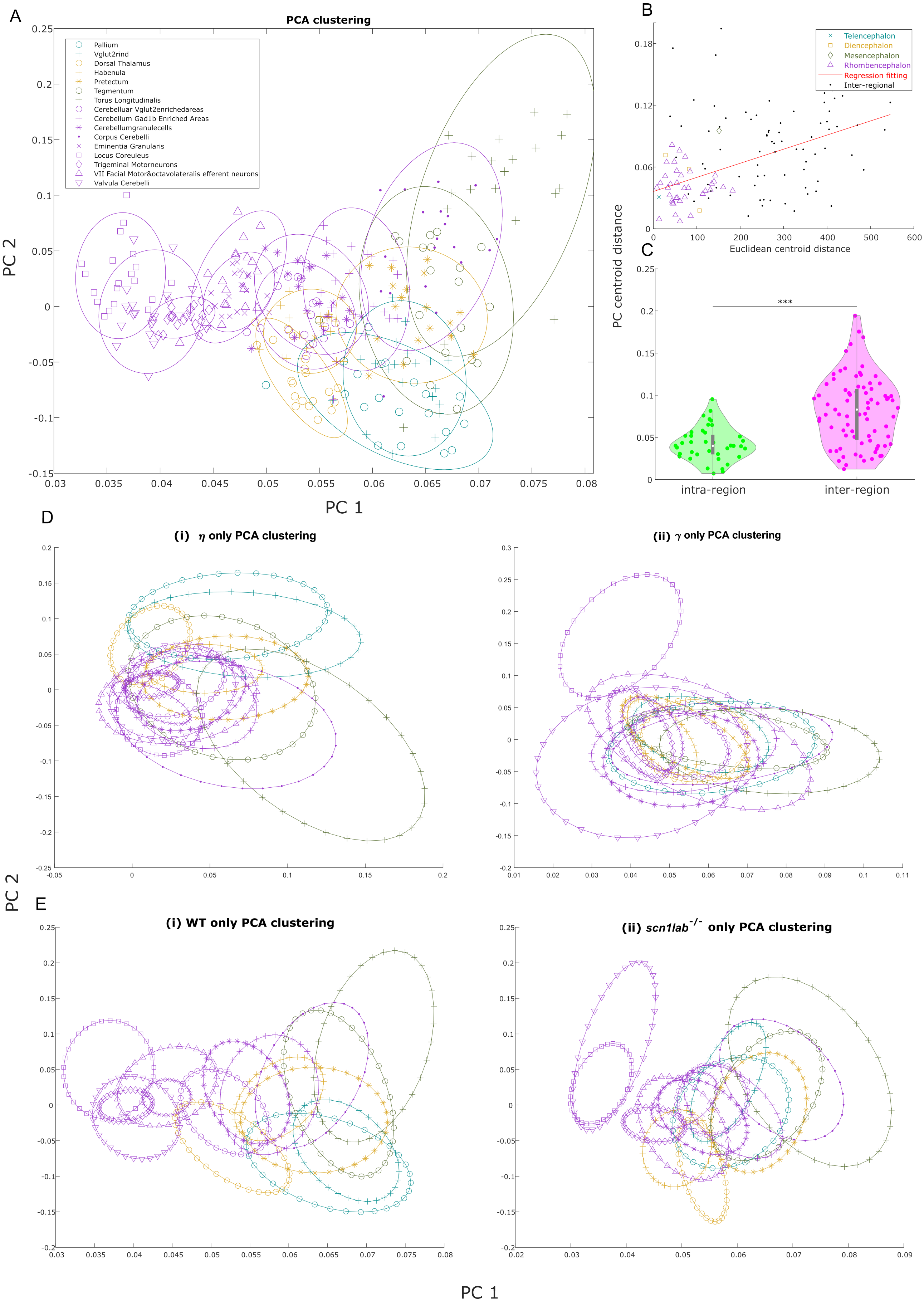
A
PCA clustering
Pallium
Vglut2rind
Dorsal Thalamus
Habenula
Pretectum
Tegmentum
Torus Longitudinalis
Cerebelluar Vglut2enrichedareas
Cerebellum Gad1b Enriched Areas
Cerebellumgranulecells
Corpus Cerebelli
Eminentia Granularis
Locus Coreuleus
Trigeminal Motorneurons
VII Facial Motor&octavolateralis efferent neurons
Valvula Cerebelli
PC 2
PC 1
B
Telencephalon
Diencephalon
Mesencephalon
Rhombencephalon
Regression fitting
Inter-regional
Euclidean centroid distance
PC centroid distance
C
***
intra-region
inter-region
D
(i) η only PCA clustering
(ii) γ only PCA clustering
E
(i) WT only PCA clustering
(ii) scn1lab-/- only PCA clustering
PC 2
PC 1

Figure S6

#### Figure S6. PCA analysis on best-fit parameters

(A) Best-fit model parameters were used as features for principal component analysis (PCA) to cluster brain subregions across both genotypes. Each circle represents a subregion, derived from 16 simulations fitted to data from all fish ($n = 35$) across 20 PTZ stages. Color indicates the parent brain region: telencephalon, diencephalon, mesencephalon, or rhombencephalon.
(B) Subregions belonging to the same anatomical region show significantly shorter pairwise distances in PCA space compared to those from different regions (Mann—Whitney U test, p $<$ 0.001; statistical data not shown). Each dot represents the principal component (PC) distance between two subregions, color-coded by intra-regional (colored) or inter-regional (black) comparison .
(C) PC distances positively correlate with physical distances between subregions. This spatial pattern supports the notion that physically proximate subregions tend to share similar wiring principles and co-localize in PCA space.
(D) Contribution of individual parameters $\eta$ and $\gamma$ to regional clustering. (i) PCA using $\eta$ alone retained partial anatomical segregation, with major brain regions loosely clustered. (ii) PCA using $\gamma$ alone yielded poor separation too, with widespread intermixing of subregions. Neither parameter alone recapitulated the anatomical distinctions observed in the full model (A).
(E) Genotype-specific PCA results. (i) In WT fish, subregions remained well-separated and spatially coherent, resembling the full dataset (A). (ii) In *scn1lab*$^{-/-}$ mutants, regional boundaries were less distinct, with widespread intermixing except for the valvula cerebelli and locus coeruleus. The anatomical clustering evident in WT was largely disrupted in the mutant genotype.

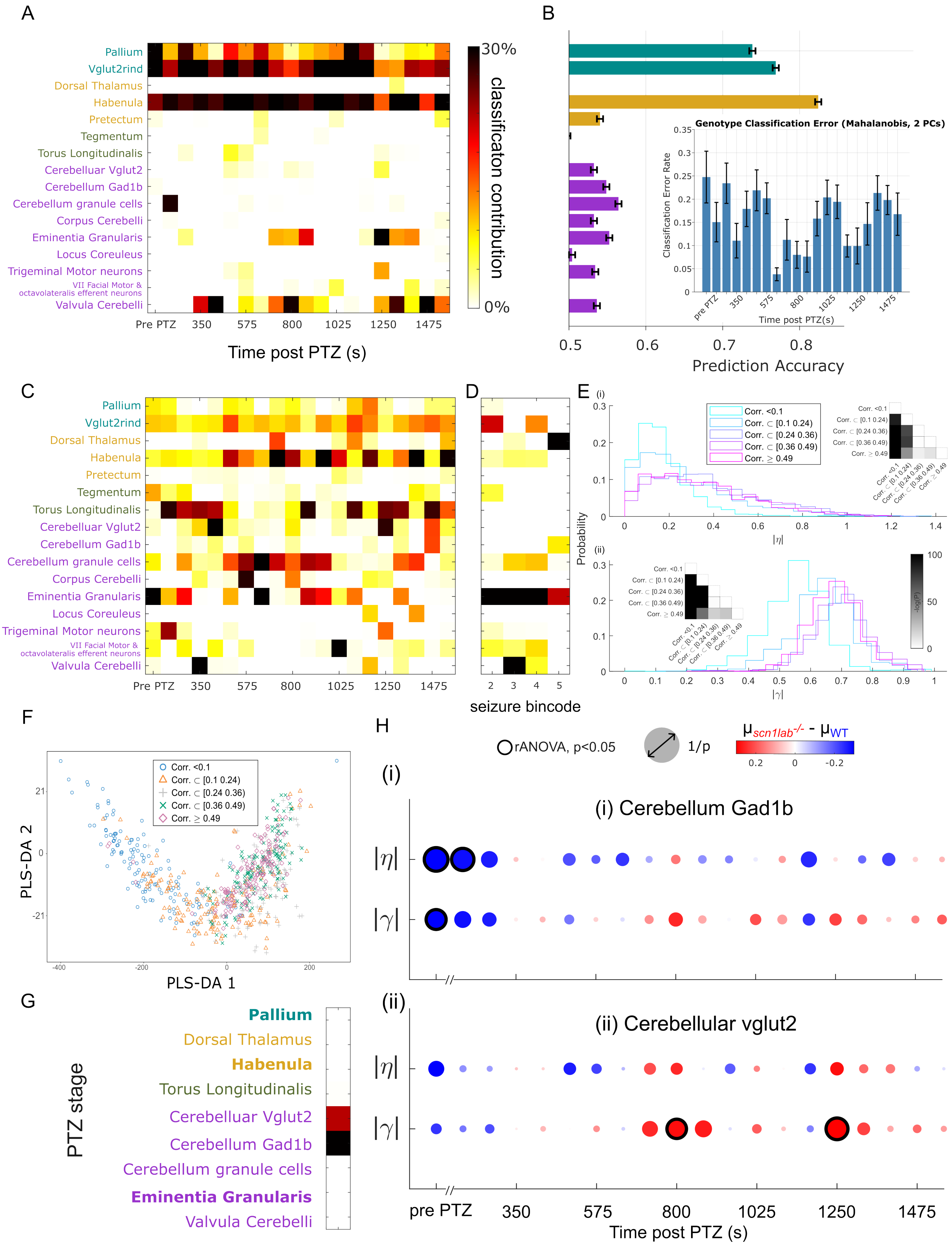
A
Pallium
Vglut2rind
Dorsal Thalamus
Habenula
Pretectum
Tegmentum
Torus Longitudinalis
Cerebelluar Vglut2
Cerebellum Gad1b
Cerebellum granule cells
Corpus Cerebelli
Eminentia Granularis
Locus Coreuleus
Trigeminal Motor neurons
VII Facial Motor & octavolateralis efferent neurons
Valvula Cerebelli
Pre PTZ 350 575 800 1025 1250 1475
Time post PTZ (s)
30%
classificaton contribution
0%
B
Genotype Classification Error (Mahalanobis, 2 PCs)
Classification Error Rate
Time post PTZ(s)
0.5 0.6 0.7 0.8
Prediction Accuracy
C
D
seizure bincode
E
Corr. <0.1
Corr. ⊂ [0.1 0.24)
Corr. ⊂ [0.24 0.36)
Corr. ⊂ [0.36 0.49)
Corr. ≥ 0.49
Probability
|η|
|γ|
-log(p)
F
PLS-DA 1
PLS-DA 2
G
PTZ stage
Pallium
Dorsal Thalamus
Habenula
Torus Longitudinalis
Cerebelluar Vglut2
Cerebellum Gad1b
Cerebellum granule cells
Eminentia Granularis
Valvula Cerebelli
H
rANOVA, p<0.05
1/p
μscn1lab-/- - μWT
(i) Cerebellum Gad1b
(ii) Cerebellular vglut2
|η|
|γ|
pre PTZ 350 575 800 1025 1250 1475
Time post PTZ (s)

Figure S7

#### Figure S7. PLS-DA analysis on subregional fingerprints

(A) Partial least squares discriminant analysis (PLS-DA) effectively classified zebrafish genotypes using predictor and response loadings (see Figure 7C). A contribution grid based on the full regional fingerprint represents 16 subregions; darker colors indicate stronger influence on seizure prediction.
(B) Classification accuracy of individual subregions alone at pre-PTZ. Pallium, *vglut2*-expressing regions, and habenula show significantly higher accuracy than other regions. Inset shows overall classification error across PTZ stages using the full fingerprints.
(C) PLS-DA on seizure counts, with a threshold of 5.5 separating 2 bins, low ($<$ 5.5) and high ($>$ 5.5) severity groups. Subregion contributions to seizure prediction are shown.
(D) GNM analysis at pre-PTZ reveals regional contributions across seizure count bin sizes. Eminentia granularis contributes most to seizure classification ($>$ 40%), alongside regions identified in (A). Darker colors denote higher predictive weight.
(E) PLS-DA across PTZ stages using best-fit $\eta$ and $\gamma$ values from simulations. Colors indicate correlation ranges; stair blocks show pairwise comparisons. Black infill marks significant differences ($p < 0.05$, two-sample Kolmogorov–Smirnov test). Distributions of $\eta$ and $\gamma$ shift rightward post-PTZ, indicating increased synchronization. $\eta$ and $\gamma$ during the final stages show reduced separability, consistent with Figure 7C.
(F) Each dot represents a fingerprint from one fish during a specific PTZ stage. Seizure stages are grouped into five categories based on correlation values, as in (E).
(G) Subregions contributing most to correlation-based PTZ stages are shown. Darker colors reflect stronger influence on seizure prediction.
(H) Regional $\eta$ and $\gamma$ values from cerebellum, specifically *gad1b*- and *vglut2*-expressing regions, reveal key differences. These regions were top contributors to PTZ stage classification in (G). Red indicates higher mean values in *scn1lab*$^{-/-}$ larvae, blue indicates higher values in WT. Circle size reflects confidence ($1/p$), and black rings denote statistical significance ($p < 0.05$, repeated-measures ANOVA).
